# Origins and Evolution of Seasonal Human Coronaviruses

**DOI:** 10.1101/2022.06.02.494567

**Authors:** JR Otieno, JL Cherry, DJ Spiro, MI Nelson, NS Trovao

## Abstract

Four seasonal human coronaviruses (sHCoVs) are endemic globally (229E, NL63, OC43, and HKU1), accounting for 5-30% of human respiratory infections. However, the epidemiology and evolution of these CoVs remain understudied due to their association with mild symptomatology. Using a multigene and complete genomes analysis approach, we find the evolutionary histories of sHCoVs to be more complex than previously recognized, owing to frequent recombination of CoVs, including within and between sHCoVs. Within sHCoV recombination rate was highest for 229E and OC43, and within genus highest for betaCoVs, whereas substitutions per recombination event inversely highest in NL63 and HKU1, and the alphaCoVs. Depending on the gene studied, OC43 may have ungulate, canine, or rabbit CoV ancestors, while 229E may have origins in a bat, camel or an unsampled intermediate host. HKU1 had the earliest most recent common ancestor (MRCA: 1809-1899), comprised two genetically divergent genotypes (A and B) possibly representing two independent transmission events from murine CoVs, and genotype B was genetically more diverse than all the other sHCoVs. Finally, we found shared amino acid substitutions in multiple proteins along the non-human to sHCoV host-jump branches. The complex evolution of CoVs and their frequent host switches could benefit from continued surveillance of CoVs across non-human hosts.

## Introduction

There are four seasonal human coronaviruses (sHCoVs) that are endemic globally (229E, NL63, OC43, and HKU1), accounting for 5-30% of human respiratory tract infections [1]. CoV infections primarily involve the upper respiratory tract and the gastrointestinal tract, mostly produce mild respiratory diseases, but may sometimes cause life-threatening bronchiolitis and pneumonia in infants, young children, elderly, and immunocompromised individuals [2–4]. While the sHCoVs are globally distributed with a general seasonality between December and April, the frequency of detection varies by location and time [5–11]. Differences in clinical presentation and frequencies of detection by age and patient groups have been observed between the sHCoV infections, and might be immune-mediated [1,11–13]. Repeat infections with the same sHCoV species are common, with up to a 21% re-infection rate over a six-month household study period [14, 15]. However, due to the historic association with mild symptomatology, there have been limited long-term epidemiological studies on sHCoVs compared to other seasonal respiratory viruses such as respiratory syncytial virus (RSV) and influenza virus [3, 12]. Furthermore, there are no approved antiviral agents or vaccines for the sHCoVs, with treatment being largely supportive.

The four sHCoV species are part of two CoV genera, alphacoronavirus (alphaCoV; 229E and NL63) and betacoronavirus (betaCoV; HKU1 and OC43) [16]. The sHCoVs are enveloped, non-segmented, positive-sense RNA viruses with genome sizes of 27-30 kb [17]. CoV genomes are quite diverse, in part due to a high frequency of RNA recombination [18, 19]. About two thirds of the genome comprises the open reading frame 1a/b (ORF1a/b) encoding two replicase polyproteins, with the remaining one third consisting of ORFs encoding the structural proteins hemagglutinin-esterase (HE: 1.2-1.3Kbp, in HKU1 and OC43 only), spike (S: 3.5-4.7Kbp), envelope (E: 0.2Kbp), membrane (M: 0.6-0.7Kbp) and nucleocapsid (N: 1.1-1.3Kbp), as well as several accessory proteins (Figure 1) [20]. The spike protein responsible for the characteristic crown-like appearance of these viruses under electron microscopy has utility in receptor binding and viral entry, and is also the most variable at the nucleotide and amino acid levels of the coronaviruses ORFs [21]. The envelope and membrane proteins have a role in the CoV assembly and determining the shape of the viral envelope while the nucleocapsid protein has a protective role through packaging of the viral genomic RNA into ribonucleoprotein complexes called nucleocapsids, ensuring timely replication and reliable transmission [22]. These four structural proteins are crucial for virus entry into host cells and subsequent replication.

**Figure 1:**
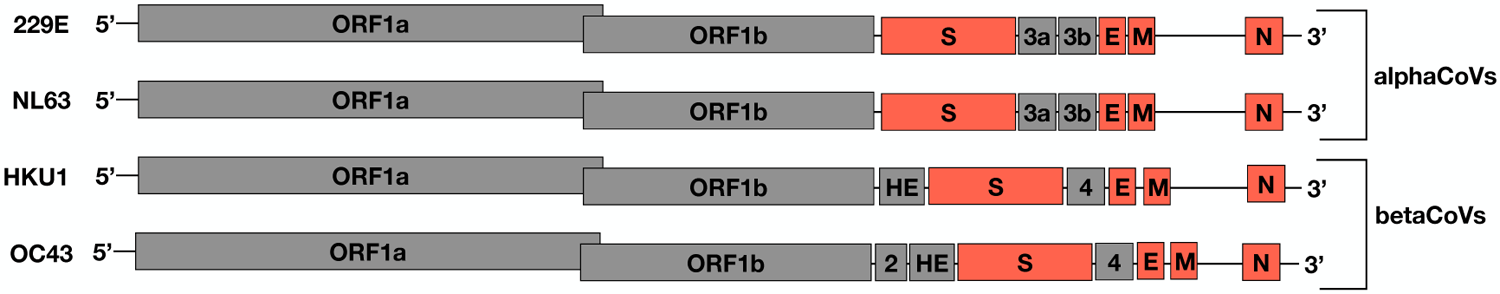
An illustration of the sHCoV genomes, not drawn to scale. In orange are the ORFs analyzed in this study.

Coronaviruses predominantly circulate among non-human hosts, with rare cross-species transmission to humans through handling of infected wild and domestic animals [23, 24]. Until 2003 and prior to the identification of the severe acute respiratory syndrome coronavirus 1 (SARS-CoV-1) [25, 26] as the causative agent of the SARS pandemic of 2002–2003 [27], only 229E and OC43, isolated nearly 60 years ago [28–30], were known to infect humans. Not long after the identification of SARS-CoV-1, NL63 and HKU1 were identified in 2004 and 2005, respectively [31, 32]. It is thought that all sHCoVs originate from bat CoVs, with some transmitted directly to humans (NL63) and others indirectly transmitted through intermediate hosts (229E, HKU1 and OC43) [33–35]. The presumptive ancestor of the OC43 is the bovine CoV, with which it has been estimated to share ∼97% genome identity [36, 37], while 229E is thought to have emerged from a camelid CoV and sharing ∼91-93% genome identity [38–40]. Lastly, HKU1 is thought to have emerged from a murine CoV [41]. Understanding the zoonotic origins of sHCoVs not only aids in directing surveillance efforts for future potential emergence of CoVs, but also highlights cultural practices at the human and non-human interface that can help in designing prevention and treatment strategies.

While there have been few epidemiological studies on sHCoVs, evolutionary studies on the sHCoVs are even more limited. As we ponder the future trajectory of the SARS-CoV-2 virus, the endemic sHCoVs provide a unique window into CoV emergence and evolutionary history in the human population. In this study, we explored the evolutionary history of the sHCoVs using a dataset of 855 sequences from GenBank, and examined the timing of emergence, zoonotic origins, genetic diversity, recombination patterns and rates, and potential adaptive protein changes. We used a multigene analysis approach that provides insight into the evolutionary histories of the spike, nucleocapsid, membrane and envelope ORFs of these CoVs. We find that the origins of the sHCoVs are much more complex and uncertain than previously thought, with our evolutionary reconstructions suggesting that different ORFs in a single sHCoV might have distinct origins. The complex and uncertain evolutionary histories of sHCoVs may be in part due to recombination and undersampling particularly of non-human hosts, and a fuller understanding will be achieved with more sequence data and re-analysis.

## Materials and Methods

### Datasets and subsampling

Dataset D1. We retrieved all coronavirus sequences available in GenBank as of 6^th^ June 2020, approximately 65,000 sequences, comprising at least 27 human and non-human hosts. This dataset was cleaned to remove sequences obtained from laboratory hosts, duplicates of the same isolate, samples that had undergone more than 10 passages, contaminants (e.g., primers and vaccines), sequences shorter than 50% of the respective protein or full genome length, and sequences for which no collection date could be reliably obtained either through GenBank metadata or publications. The dataset was subsequently restricted to the alphaCoV and betaCoV genera (to which the four sHCoV species belong), and focused the analyses on the envelope, membrane, nucleocapsid and spike proteins and whole genome sequences (WGS) (*n*=8032 sequences, Supplementary Figure 1).

Dataset D2. There was an obvious sampling bias in the dataset D1 by host and time (Supplementary Figure 1), and these biases are known to impact phylodynamic inferences [42, 43]. To address this, Maximum Likelihood (ML) trees (see section on Maximum Likelihood (ML) trees and temporal signal) were generated with the D1 dataset for each structural protein and WGS, and then downsampled each non-human CoV to ∼ 30-40 sequences per host-clade (since some CoVs from the same host were represented by multiple clades) using the phylogenetic diversity analyzer software (PDA) that selects a subset of sequences that comprises the maximal phylogenetic diversity [44]. The earliest collected CoV sequence(s) for each host-clade were always retained in the PDA subsampled datasets. (*n*=2382 sequences).

Dataset D3. For computational efficiency, dataset D2 was further downsampled to a clade of coronaviruses with bootstrap support >0.7 and a reasonably sufficient number of the non-human hosts (Supplementary Figure 2). In accordance with previous reports and observed preliminary clustering patterns on ML trees in this analysis, the HKU1 sequences were partitioned into two genotypes A and B [8, 45]. (*n*=1729 sequences).

Dataset D4. To avoid the potential impact of the much larger number sHCoV sequences in comparison with their respective non-human CoVs ancestors, the 229E and NL63 sequences were downsampled to an equal number of sequences as those of the 229E-related camelid CoVs [38–40]. For betaCoVs, the sHCoVs OC43 and HKU1 sequences were downsampled to an equal number as those of murine CoVs that were inferred to be closest phylogenetically [41]. Similar to Dataset D2, PDA was used for the downsampling and the earliest isolated sHCoV sequence(s) for each species were always included in the subsampled datasets. (*n*=855 sequences, <u>Supplementary File 1</u>). Multiple sequence alignments were built using MAFFT v7.475 [46] and thereafter edited and cleaned manually using AliView v1.26 [47].

### Maximum Likelihood (ML) trees and temporal signal

For each dataset described above, an ML tree was generated using IQ-TREE [conda versions 2.0.3-2.1.4] [48] allowing the software to determine the best nucleotide substitution model [49] and using ultrafast bootstrap [50] approximation to estimate branch support. In order to visually examine the degree of temporal signal or accumulation of divergence in the datasets over the sampling time interval, the exploratory linear regression approach implemented in TempEst v1.5.3 [51] was employed. The root-to-tip divergences as a function of sampling time according to a rooting that maximizes the Pearson product moment correlation coefficient was plotted in TempEst. Outlier sequences that were either not divergent enough or too divergent were removed from the datasets. The new datasets were used to generate new ML trees that were heuristically time-transformed using TempEst and subsequently used as starting trees to reduce the burn-in of the Markov Chain Monte Carlo (MCMC) phylodynamic analyses (see section on Phylodynamics and host-jump analysis).

### Comparative genomics

With dataset D4 and using MEGAX [52], we calculated the mean sequence divergence over all sequence pairs for each CoV species. We used the maximum composite likelihood method [53], assumed a heterogenous substitution pattern among lineages, and modelled evolutionary rates among sites using the gamma distribution with four rate categories.

The non-human CoVs selected for this formed a clade with the sHCoVs and were either paraphyletic to the sHCoVs or a sister group with bootstrap support >0.7. However, when the non-human CoVs that shared an MRCA with sHCoVs, rather than the ones in the more distant past, were represented by a single sequence, the mean pairwise genetic distance could not be calculated on the single sequence.

### Recombination analysis

Coronaviruses are known to frequently recombine [54]. We used ClonalFrameML v1.12 and dataset D4 to estimate the ratio of the rate of recombination to the rate of point mutation (*ρ/θ*, also referred to as *R/theta* or *ε*), the ratio of effects of recombination to point mutation (*r/m*), and the number of substitutions per recombination event (*δν*). The *R/theta*, *r/m* and *δν* values were estimated not only within the alphaCoVs and betaCoVs, but also within sHCoV species. To perform comparisons of these estimates between sHCoV species, we used subsets of dataset D4 that (i) had one sHCoV species and all other non-human CoVs [to estimate inter-host recombination rates], and (ii) had one sHCoV species [to estimate intra-species recombination rates]. However, since ClonalFrameML was designed to be used on bacterial full genome datasets, this analysis was only performed on WGS (29-32 Kbp) and the spike protein (4.1-4.7 Kbp).

Dataset D5. Recombination can distort phylogenetic tree inference and interpretation [55–57]. Using RDP4 [58], recombinant sequence regions in dataset D4 were iteratively removed until no more recombination signals were detected. The software’s default settings were used except for specifying that the sequences were linear and adding the 3SEQ method to the five default methods (RDP, GENECONV, MaxChi, Bootscan and SisScan). Recombination signals were considered present if they were detected by three of more methods. Sequences that lost more than 50% of the respective ORF or WGS lengths as a result of the removal of the recombinant regions were discarded. (*n*=783 sequences).

Dataset D6. To test for recombination between sHCoVs from different genera, all the sequences from dataset D4 for WGS and each ORF were combined into single datasets and the non-sHCoV sequences discarded. Recombination analysis was performed on dataset D6 with RDP as described above. (*n*=346 sequences).

### Phylodynamics and host-jump analysis

We used the BEAST v1.10.4 package [59, 60] and dataset D5 to estimate the timing of the emergence of each sHCoV species, evolutionary rates, and zoonotic origins. Conditioning on the host of the taxon, the host transition history was modelled as a non-reversible continuous time markov chain (CTMC) process while reconstructing the unobserved hosts at the ancestral nodes in each tree of the posterior distribution [61]. For sequences with incomplete dates, we incorporated an uncertainty in the sampling date. An HKY+G+I nucleotide substitution model [62] was specified together with a Skyline coalescent tree prior [63] and an uncorrelated relaxed clock model [64]. The MCMC sampling in BEAST was undertaken for at least 700 million steps for each run, with each dataset undergoing at least three independent runs. We combined the log and tree files using LogCombiner until the effective sampling sizes (ESS) were >200 as assessed by Tracer v1.7.1 [65]. The posterior tree distributions were summarized and annotated with TreeAnnotator, after discarding at least 10% of the trees as chain burn-in. The summarized maximum clade credibility (MCC) trees were visualized using FigTree v1.4.4 and ggtree [66].

We also estimated the number of host transitions (Markov jumps) in order to have a complete summary of the host transmission processes [67]. (GitHub location for XML files: to be updated upon final submission)

### Selection analysis

Using the Datamonkey server (https://www.datamonkey.org), we tested for positive selection along the host-jump branches leading to the sHCoVs using two separate methods: (i) the adaptive Branch-Site Random Effects Likelihood (aBSREL) method [68], and (ii) the Branch-site Unrestricted Statistical Test for Episodic Diversification (BUSTED) method [69]. Both aBSREL and BUSTED estimate the ratio of nonsynonymous to synonymous substitutions (ω), determine the optimal number of ω rate classes, and estimates the proportion of sites belonging to each ω rate class. For this analysis, we used BEAST MCC trees with recombination-free tree topologies (from dataset D5) and sequence alignments from dataset D4 with recombinant regions intact so as not to miss potential positive selection signals from such regions.

### Amino acid substitution analysis

We used BEAST MCC trees (from dataset D5) and sequence alignments from dataset D4 to perform amino acid (AA) ancestral reconstruction [70] of historical CoVs in order to elucidate the AA changes that might have characterized the inter-host transmission of non-human CoVs into humans, as well as the evolution and adaptation of the sHCoVs within the human host. The ancestrally reconstructed AA changes were also compared with the AA changes characterizing the SARS-CoV-2 Wuhan-Hu-1 or a variant of concern (VOC). The dataset D4 sequences were aligned against the Wuhan-Hu-1 SARS-CoV-2 reference genome (NC_045512.2), after which the reference sequence was removed prior to the ancestral reconstruction and AA analysis. Where there was an AA insertion in the sHCoVs relative to the Wuhan-Hu-1 SARS-CoV-2 reference, we used the X.Y positional notation where X is the reference genome position and Y is the n^th^ sHCoV AA insertion, i.e. the 2 in AA position 8.2 indicates the second AA insertion at position 8 relative to the Wuhan-Hu-1 SARS-CoV-2 reference genome.

## Results

### Zoonotic origins of the seasonal human coronaviruses

To investigate the zoonotic origins of the four sHCoV species, we generated MCC trees from the WGS, and the spike, envelope, membrane and nucleocapsid ORFs.

For 229E, MCC trees generated from the WGS and the four ORFs had different topologies arising from recombination events deep in the viral evolutionary history. Bats are the ancestral host on each tree, but the phylogenetic relationships between sHCoV 229E and clades associated with other CoV host species differ (Figure 2). The 229E and camelid CoVs formed a clade with very high branch support (posterior probability [PP] = 1) on the spike protein and WGS MCC trees. However, 229E was more closely related to bat CoVs than to camelid CoVs on trees inferred for the nucleocapsid, membrane, and envelope proteins. Analysis of the CoVs inter-host transmission process (markov jumps [MJs]) for the envelope, membrane and nucleocapsid estimated a 98-99% probability that the sHCoV 229E arose directly from a bat CoV (Table 1). For the spike protein and WGS, there was a 73% and 65% probability that 229E arose directly from a bat CoV, and a 27% and 34% probability that 229E originated from a camelid CoV, respectively. As camelid CoVs did not seem to have the higher probability of being ancestral to sHCoV 229E, we looked at the estimated origins of these camelid CoVs. It was estimated for the spike and WGS that there was a 61% and 35% probability, respectively, that the camelid CoVs arose from the sHCoV 229E. Taken together, there is a high degree of uncertainty as to whether camelids were intermediate hosts for the transmission of a bat CoV into humans or whether transmission occurred directly from a bat, or potentially via an unsampled intermediate host, given the long branch lengths. This analysis also considers the possibility of a reverse zoonosis of the human 229E CoVs into camelids.

**Figure 2:**
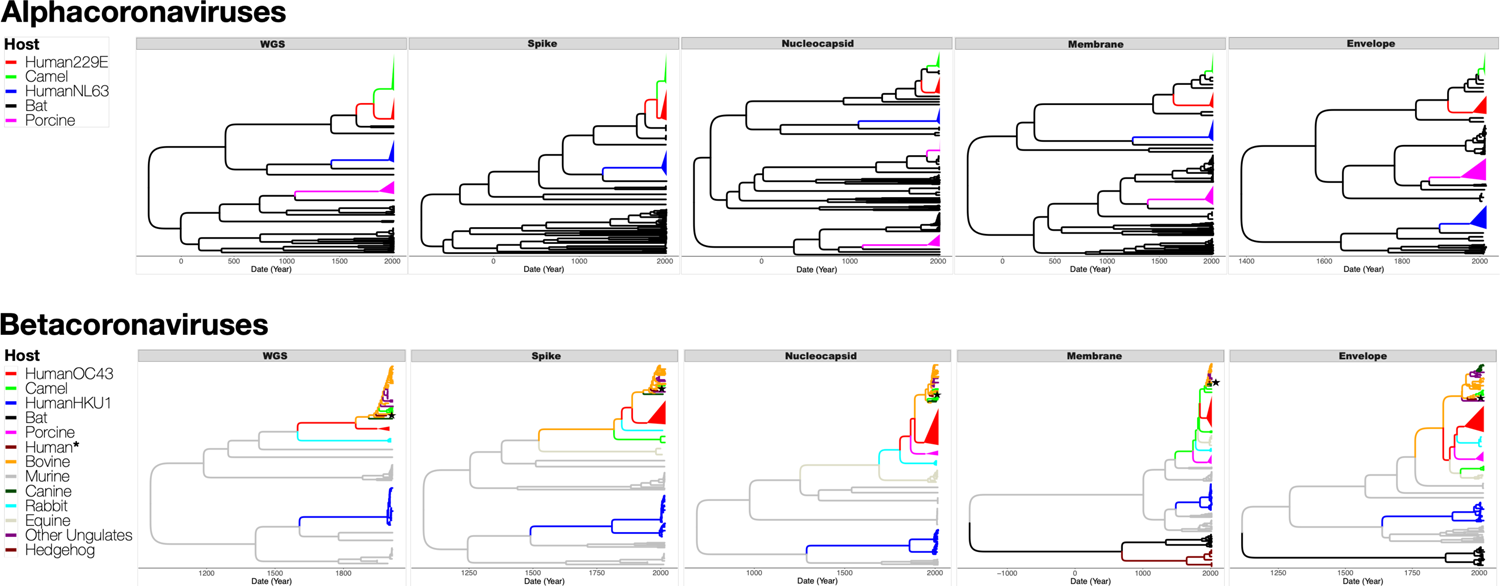
Maximum clade credibility (MCC) trees inferred from dataset D5 for full genomes (WGS), and the spike, nucleocapsid, membrane and envelope proteins, with the branches color-coded by the inferred coronavirus host. The upper panel shows MCC trees from alphacoronaviruses while the lower panel shows MCC trees from betacoronaviruses. Human, camel and porcine coronavirus clades have been collapsed to increase readability. Human* is a lone human CoV (FJ415324) that clusters with ungulate and canine CoVs. Individual and more detailed MCC trees can be found in <u>Supplementary File 2</u>.

**Table 1:**
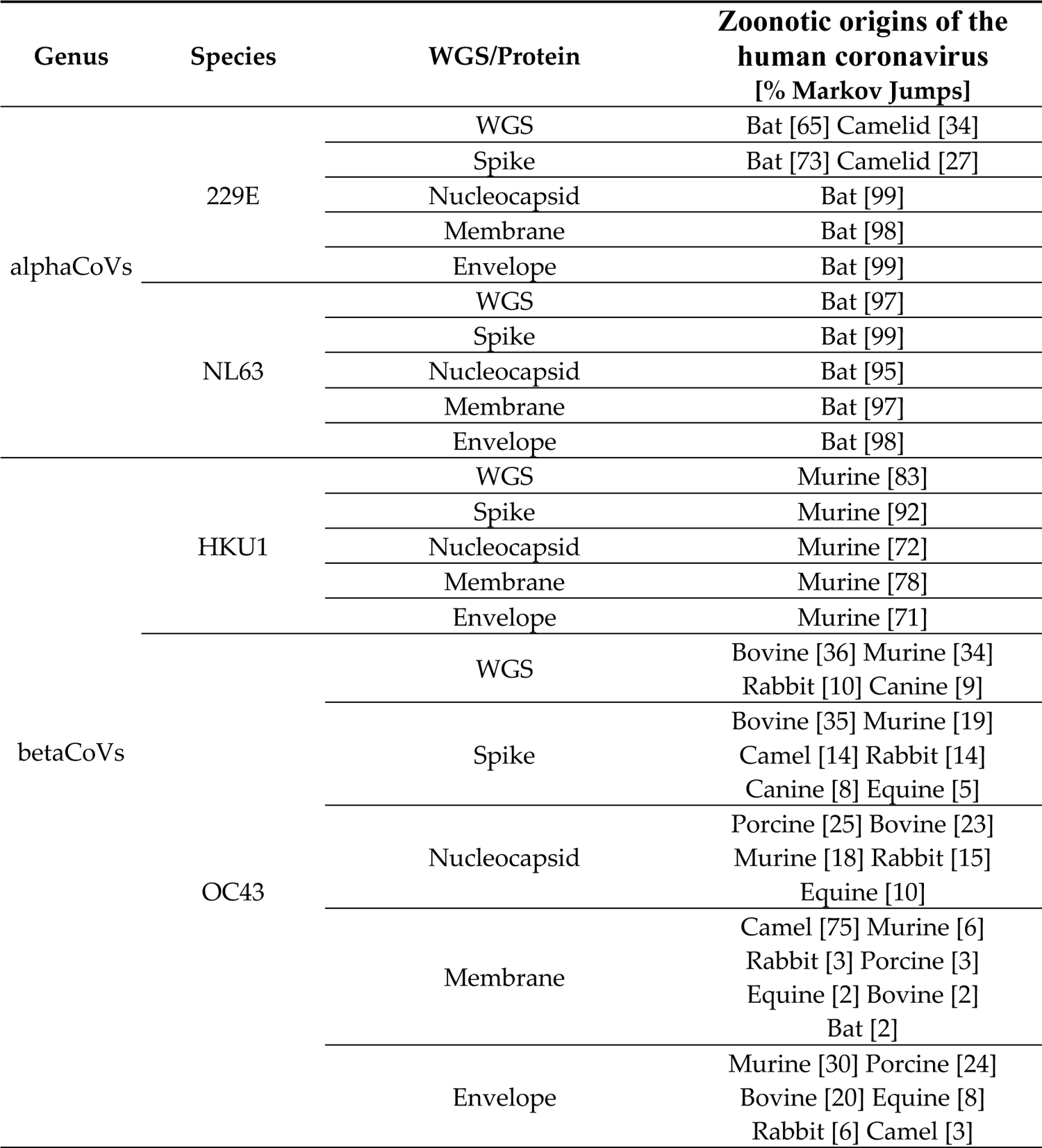
The estimated zoonotic origins of the four species of human seasonal coronaviruses inferred from BEAST Markov Jumps (MJs) for whole genomes and the envelope, membrane, nucleocapsid and spike proteins. For 229E and OC43, we included the percentage of MJs of the presumptive ancestors, other potential ancestors with a percentage higher than the presumptive ancestor, or CoVs whose cumulative percentage of MJs were ≥70%.

The zoonotic origins of the sHCoVs NL63 and HKU1 were inferred to be bat and murine CoVs, respectively, with no apparent intermediate hosts (Figure 2 and Table 1). The two HKU1 A and B genotypes formed distinct and well-supported clades, with very long branch lengths, in all the MCC trees derived from the four structural proteins (Figure 2).

Our analysis revealed that the sHCoV OC43 was a sister clade to a large group of CoVs from ungulate hosts (included bovine, buffalo, camel, deer, waterbuck, and yak) and canines, on MCC trees derived from WGS or from the spike, nucleocapsid and membrane proteins (Figure 2). Bovine CoVs were estimated to be the ancestor to OC43 with probabilities of 36%, 35%, 23%, 2% and 20% for the WGS, spike, nucleocapsid, membrane, and envelope proteins, respectively (Table 1). The host with the highest probability of CoV origins for OC43 was a porcine CoV (25%) for the nucleocapsid, a camel CoV (75%) for the membrane, and a murine CoV (30%) for the envelope. While this analysis is consistent with a bovine CoV being the origin of the OC43 [36, 37], it broadens the range of plausible origins to include murine, rabbit, canine or other ungulate CoVs.

### Rates of evolution and emergence dates

The mean substitution rates for the four sHCoVs were similar for the WGS and the four ORFs, ranging between 2.1 x 10^-4^ and 6.4 x 10^-3^ nucleotide substitutions/site/year and with overlapping 95% highest posterior density (HPD) (Figure 3A). However, the mean substitution rates for the ORFs had wider 95% HPDs than the full genomes, which is consistent with what would be expected from whole genomes that have more genetically informative sites in the estimation of the substitution rates. Nonetheless, even with the overlapping 95% HPD, HKU1 and OC43 coronaviruses often had the highest and lowest, respectively, mean substitution rates.

**Figure 3:**
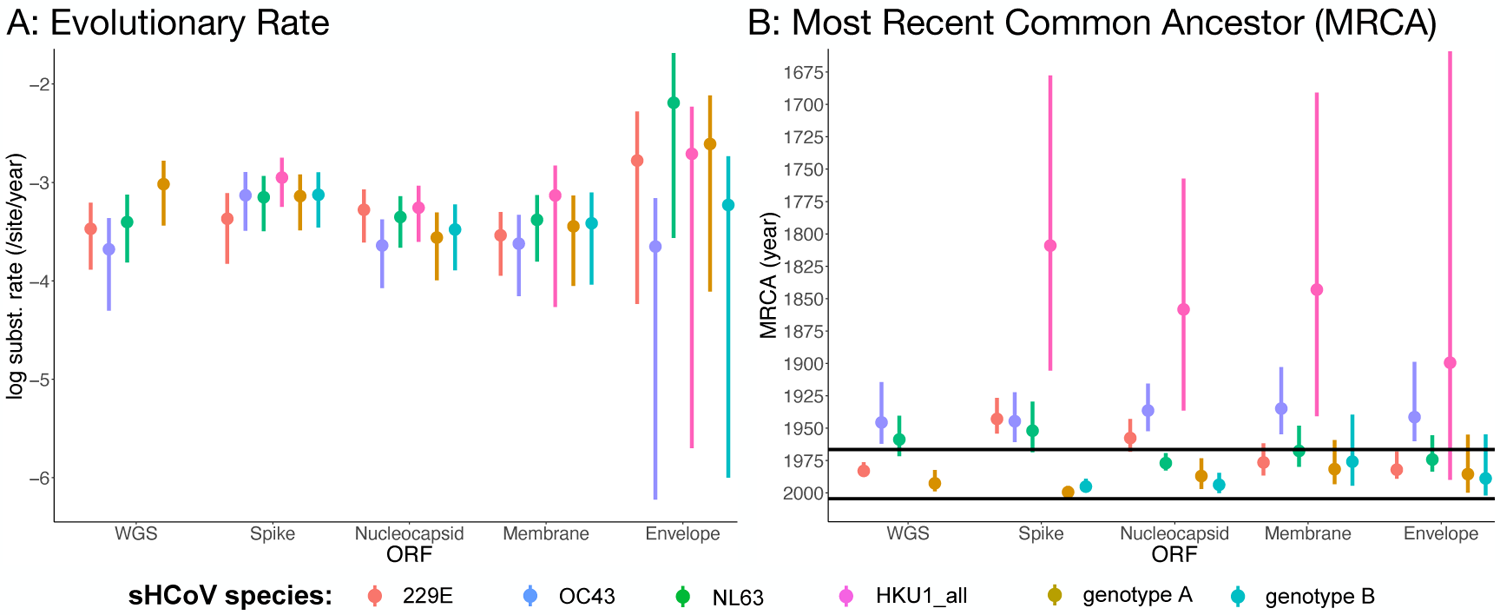
Estimates of the evolutionary rate (A) and MRCA age (B) for full genomes and four open reading frames (dataset D5) of the seasonal human coronavirus species. The black horizontal lines in (B) are the dates of first isolation for the 229E (1966), OC43 (1967), NL63 (2004) and HKU1 (2005). The WGS is missing data points for HKU1_all (collective for both genotypes) and genotype B as sequences for genotype B were all removed in the generation of recombination free WGS dataset D5.

While the dates on which the sHCoVs were first isolated are well known [28,30,32,71], we were interested in estimating their MRCAs (Figure 3B and Supplementary Figure 3). The median MRCA of the HKU1 viruses (HKU1_all) for the WGS, spike, nucleocapsid and envelope was dated between 1809 and 1899 with wide 95% HPDs, and was much older than the median MRCAs of the other sHCoVs (median: 1934-1983). However, HKU1 genotypes A and B had their median MRCAs dated much more recently between 1975 and 1999, with overlapping 95% HPDs and we could not determine which of the two HKU1 genotypes might have emerged first. It is only for the spike ORF that the median MRCAs for the 229E, OC43 and NL63 appeared to follow the sequence of their respective first isolation dates. While the spike had the earliest median age of the MRCAs for 229E, NL63 and HKU1, this ORF had the youngest median MRCA ages for OC43 and HKU1 genotypes A and B. The estimated much earlier dating of HKU1 was not expected given these CoVs are the most recently isolated of the sHCoVs [32], and raises questions whether this could be a consequence of undersampling and/or dating of two very distinct variants (genotypes A and B) that were independently transmitted into the human host.

### Recombination patterns

Recombination is a known phenomenon in CoVs, and we identified recombination signals along the evolutionary history of the alpha and beta CoVs in our datasets (Figure 4 and Supplementary File 3). With the exception of the envelope protein, recombination was detected by RDP4 in the membrane, nucleocapsid, spike, and WGS datasets. Some sHCoV sequences were characterized to be recombinants of CoVs from two or more hosts, as has similarly been shown for the SARS-CoV-2 spike protein [72]. Further inspection of the recombinant sequence pairs revealed that in addition to inter-host CoV recombination, there is recombination between human CoVs, both within and between sHCoV species (Figure 4). While intra-species recombination has previously been reported for genotypes within OC43 and HKU1 [45, 73], and between a few NL63 isolates [74], we identified recombination within 229E as well as between the sHCoV species. However, recombination between sHCoV species was only observed between species in the same genus, i.e., between 229E and NL63 and between HKU1 and OC43, and was only identified in the spike and WGS datasets. Because sHCoV co-infections are frequently identified in community and patient isolates [3, 10], recombination within and between sHCoV species might be a frequent occurrence and could increase the breadth of the sHCoV species’ variants.

**Figure 4:**
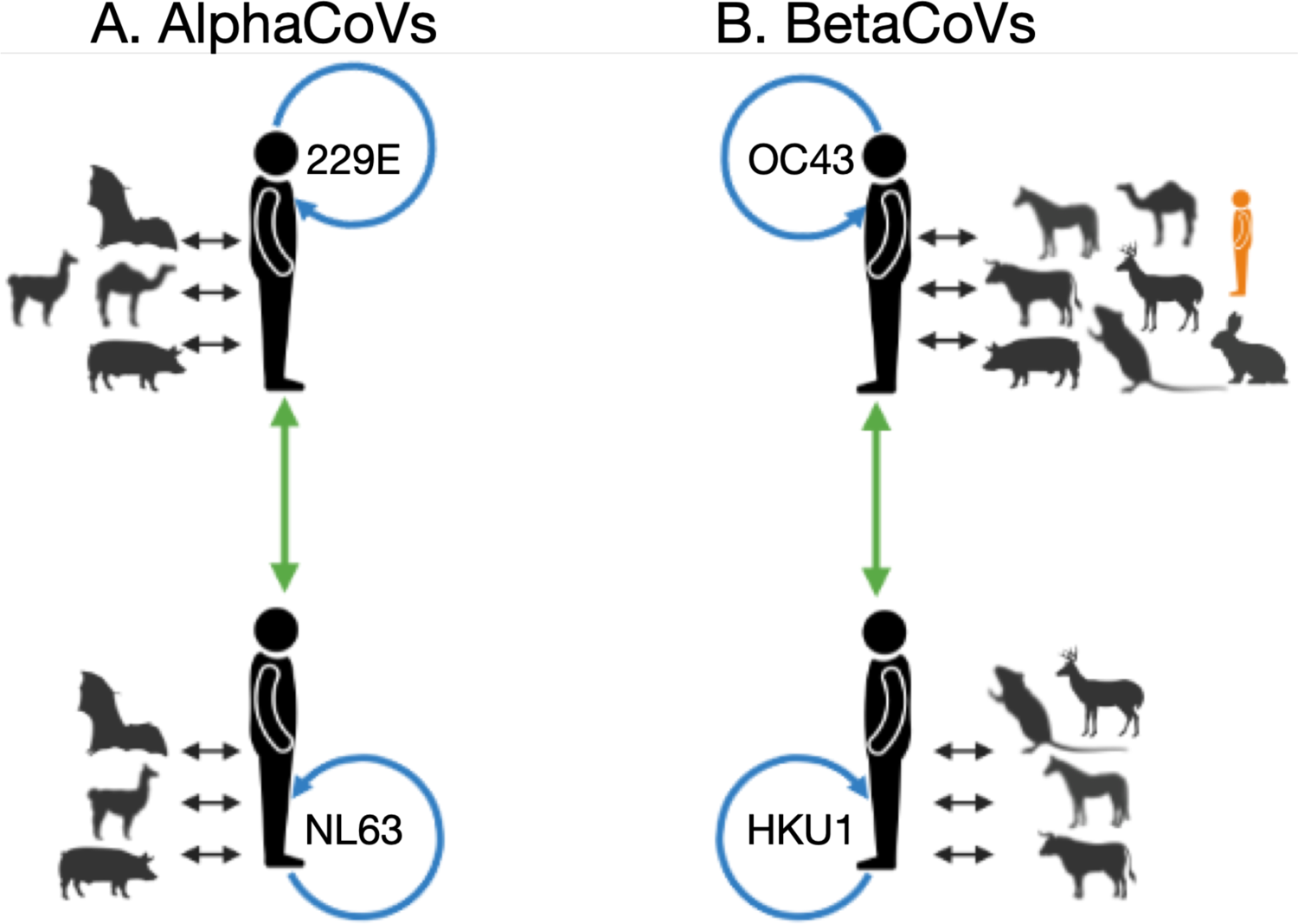
Summarized within and between host/species recombination patterns identified by RDP4, for alphaCoVs (A) and betaCoVs (B). For each sHCoV species, recombining CoVs are shown; non-human and sHCoV (black arrows), within sHCoV species (blue arrows), and between sHCoV species (green arrows). In orange is a lone human CoV (FJ415324) that clusters with ungulate and canine CoVs. Figure generated using Biorender.

### Recombination rates

The observed frequency of recombination reflects both the rate of recombination events and the effects of selection. We assessed the apparent rates of recombination among CoVs and across the genome using ClonalFrameML. We estimated that recombination happened up to two times more often than nucleotide changes due to point mutations (R/theta) in the spike protein, but 40 (betaCoVs) to 71 times (alphaCoVs) less often across the whole genome (Table 2a and 2b), indicating that the rate of recombination and/or patterns of selection might vary substantially along the genome with hotspots in certain genomic regions. The mean length of DNA imported by homologous recombination (*δ*) ranged from 209bp (betaCoVs) to 336bp (alphaCoVs) for the spike protein and 781bp (betaCoVs) to 1033bp (alphaCoVs) across the whole genome. Recombination overall was responsible for 29-42 times more substitutions than point mutation (*r/m*) in the spike, and approximately 0.8 times as many substitutions as point mutation across the whole genome. These estimates illuminate the substantial contribution of recombination in the evolution of CoVs, and particularly in the spike protein.0

**Table 2:**
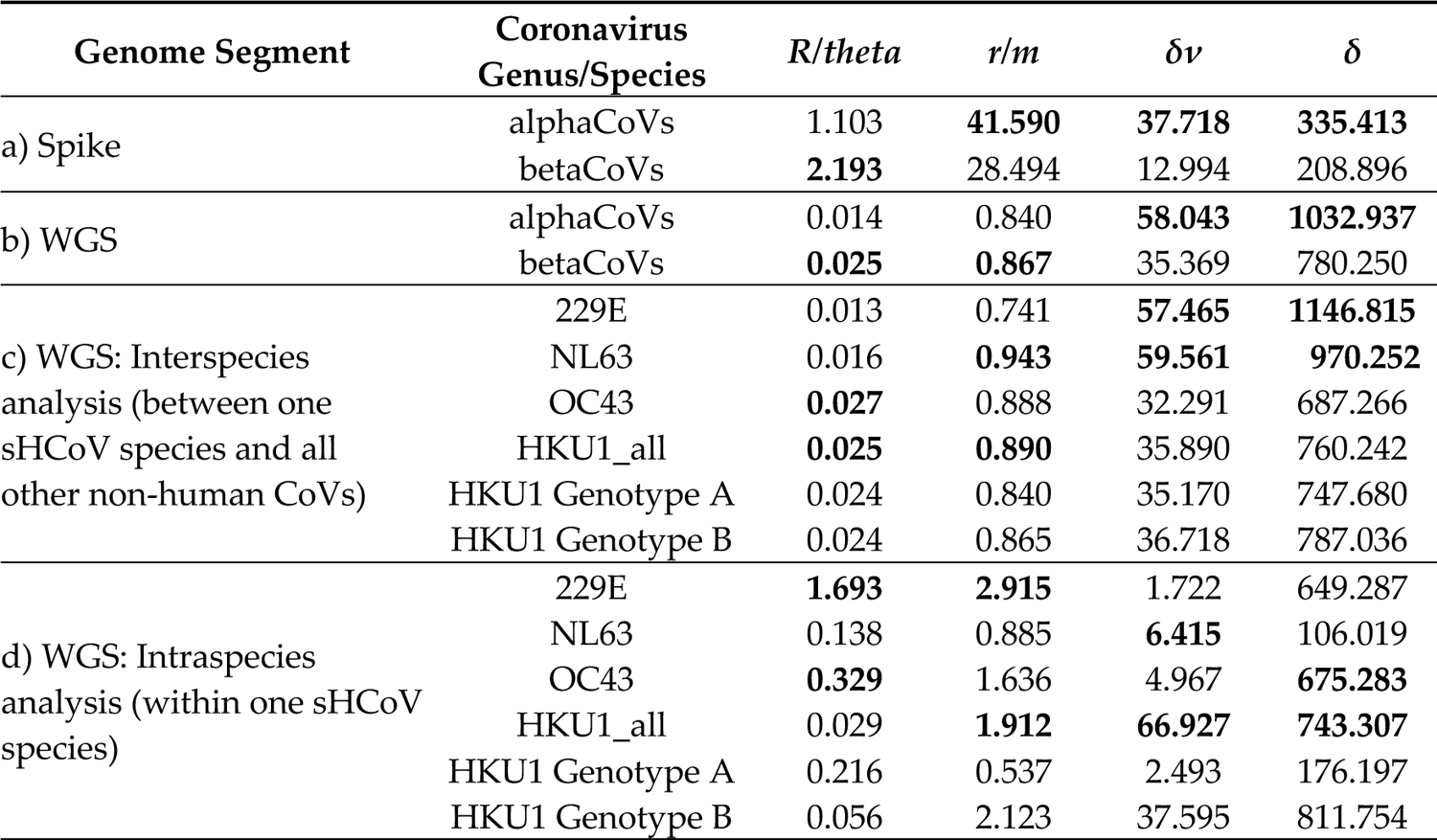
Estimates of the ratio of recombination rate to point mutation rate (*R/theta*), substitutions by recombination relative to point mutation (*r/m*), number of substitutions per recombination event (*δν*), and mean length of DNA imported by homologous recombination (*δ*) from dataset D4. For each genome segment and analysis, the highest values are shown in bold. These analyses were conducted for the two HKU1 genotypes collectively (HKU1_all) and independently.

With regard to differences in recombination rates between the alphaCoVs and betaCoVs genera (Table 2a and 2b), our estimates of *R/theta* were about two-fold higher in betaCoVs than alphaCoVs across the whole genome (0.025 vs 0.014) and within the spike protein (2.19 vs 1.10). The estimated number of substitutions introduced by each recombination event (*δν*), however, was higher in the alphaCoVs than in the betaCoVs both for the spike (38 vs 13 substitutions) and across the whole genome (59 vs 36 substitutions). These results were consistent with subset datasets comprised of one sHCoV species and all other non-human CoVs (Table 2c). These observations imply that although the rate of recombination to point mutation is higher for the betaCoVs, more substitutions are introduced per recombination event for the alphaCoVs.

Nevertheless, the distinctive patterns of *R/theta* for alphaCoVs vs betaCoVs were not consistent for the within sHCoV species recombination rate analysis (Table 2d), and instead followed the following order: HKU1<NL63<OC43<229E. This trend was reversed when considering the number of substitutions introduced by each recombination event, i.e. highest in HKU1 (67 substitutions) and lowest in 229E (2 substitutions). With regard to the HKU1 genotypes, there was a marked difference in the estimates of *R/theta* (0.2 vs 0.06) and *δν* (2 vs 37 substitutions) for genotypes A and B, respectively. The intra-genotype *R/theta* value for HKU1 genotype A was higher than that for NL63 while its corresponding *δν* value was only higher than that for 229E, depicting interesting variation in these *R/theta* and *δν* values between the two HKU1 genotypes and in comparison with the existing sHCoV species. We surmise that while sHCoV interspecies recombination rates tend to follow the species’ genera (highest for betaCoVs), intraspecies recombination rates seem to follow sHCoV first isolation dates (highest for 229E and OC43) which would allow for sufficient diversity to be accumulated and thereby the recombination signal to be detectable.

### Pairwise diversity

The estimated mean pairwise genetic diversity for each sHCoV is presented in Table 3. Genome-wide, 229E was estimated to be the least genetically diverse of the sHCoVs with HKU1 being the most genetically diverse, i.e. 229E<OC43<NL63<HKU1. This order of genetic diversity was similar to the order from ClonalFrameML’s number of substitutions introduced by each recombination event (*δν*). HKU1 genotype B was independently more genetically diverse than 229E, NL63 and HKU1, while HKU1 genotype A was the least diverse. Of the four ORFs analyzed, the spike ORF was the most genetically diverse for the sHCoVs as well as for the CoVs from non-human hosts (Table 3 and Supplementary Table 1).

**Table 3:**
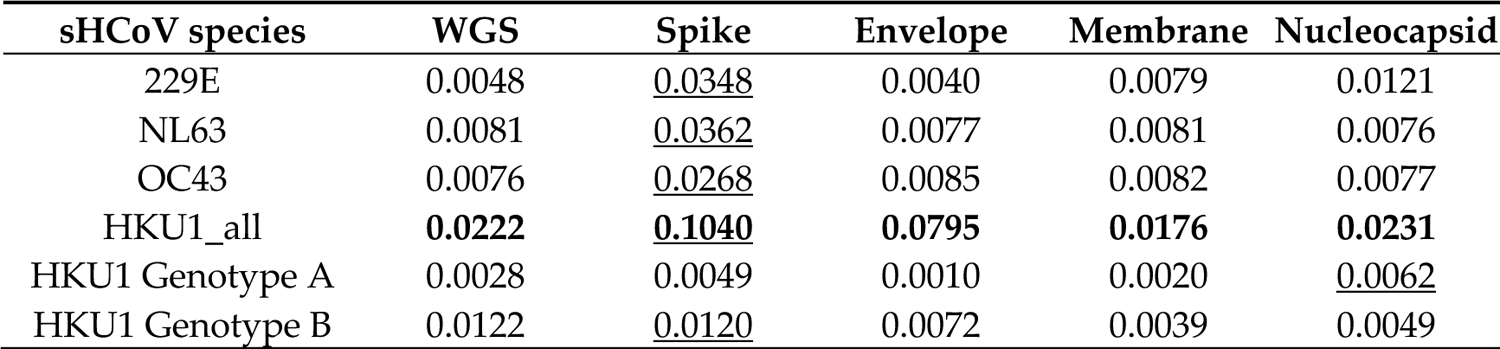
Mean pairwise genetic distances for each sHCoV species derived from WGS and four ORFs from dataset D4. In bold is the highest pairwise distance for each ORF/WGS, and underlined is the highest pair-wise distance for each sHCoV species.

Bat and murine CoVs that were sister groups to the sHCoVs were at least 7 to 9 times more diverse than the sHCoVs from the whole genome estimates (Supplementary Table 1). In fact, murine CoVs were 62 and 14 times more diverse than the HKU1 genotypes A and B, respectively. Bat CoVs were the most diverse of the non-human CoVs while the ungulates-canines CoVs that share an MRCA with OC43 were the least diverse. Lastly, camelid CoVs that share an MRCA with 229E were less diverse compared to the human 229E CoVs.

### Selection analysis

We hypothesized that there might be positive selection on the CoV host-jump branches into humans for the envelope, membrane, nucleocapsid and spike proteins due to their role in virus entry into host cells and subsequent replication. BUSTED did not infer any of the branches leading to the sHCoVs to be under positive selection (Table 4). On the other hand, aBSREL inferred the branches leading to the emergence of NL63, HKU1_all, and HKU1 genotype B to be under positive selection for the nucleocapsid protein. However, the high ω values estimated for a minority of codons seem unrealistically high, and likely reflect misalignment of a minority of codons rather than adaptation.

**Table 4:**
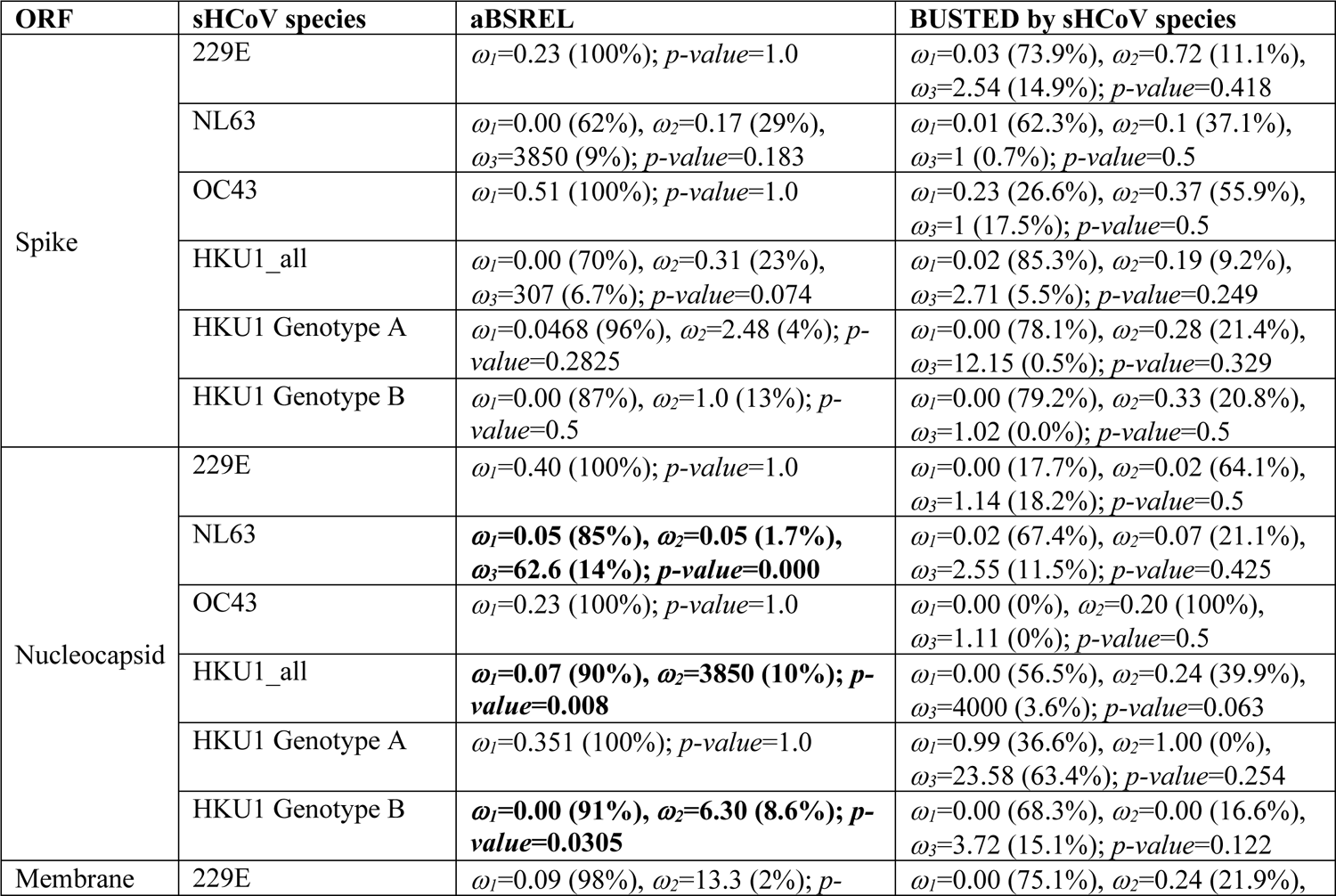
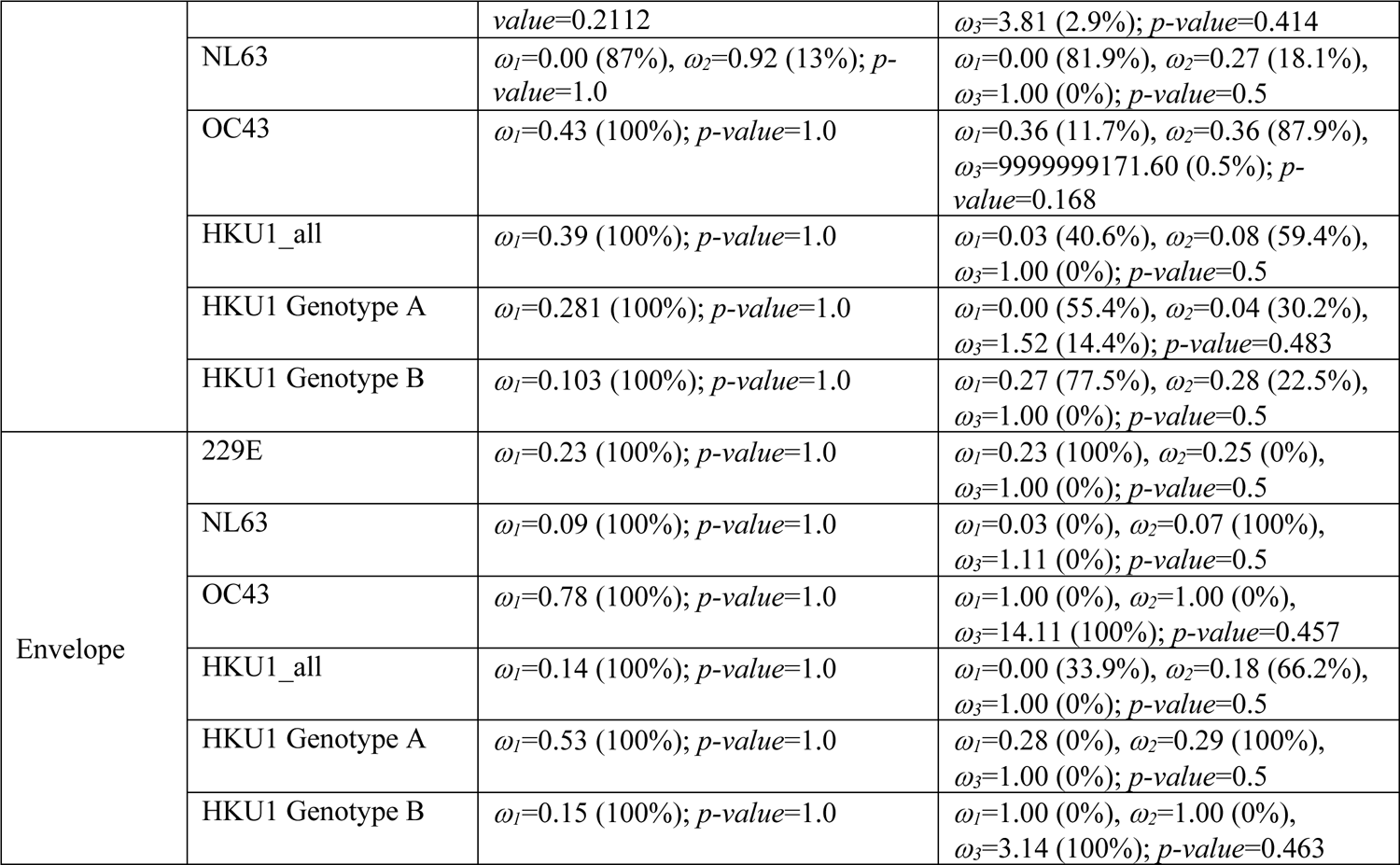
Test for positive selection in the emergence of sHCoV species from datasets D4 and D5. In bold are the host-jump branches where positive selection was inferred using the likelihood ratio test at a threshold of *p*≤0.05. Also shown are *ω* (ratio of nonsynonymous to synonymous substitutions) and the proportion of sites in each rate category.

### Amino acid substitutions

Amino acid (AA) ancestral reconstruction revealed that 29-57%, 29-35%, 16-26% and 19-48% of the AA alignment positions of the spike, nucleocapsid, membrane and envelope, respectively, had at least one AA substitution across the genealogy of the sHCoVs (Supplementary File 4). In Figure 5 (and Supplementary Figure 4) we show the number of the inferred AA changes associated with the sHCoVs in the envelope, membrane, nucleocapsid and spike proteins. While AA changes were inferred to occur throughout the spike protein, an elevated number of changes occurred within the N-terminal domain (NTD) and the receptor binding domain (RBD).

**Figure 5:**
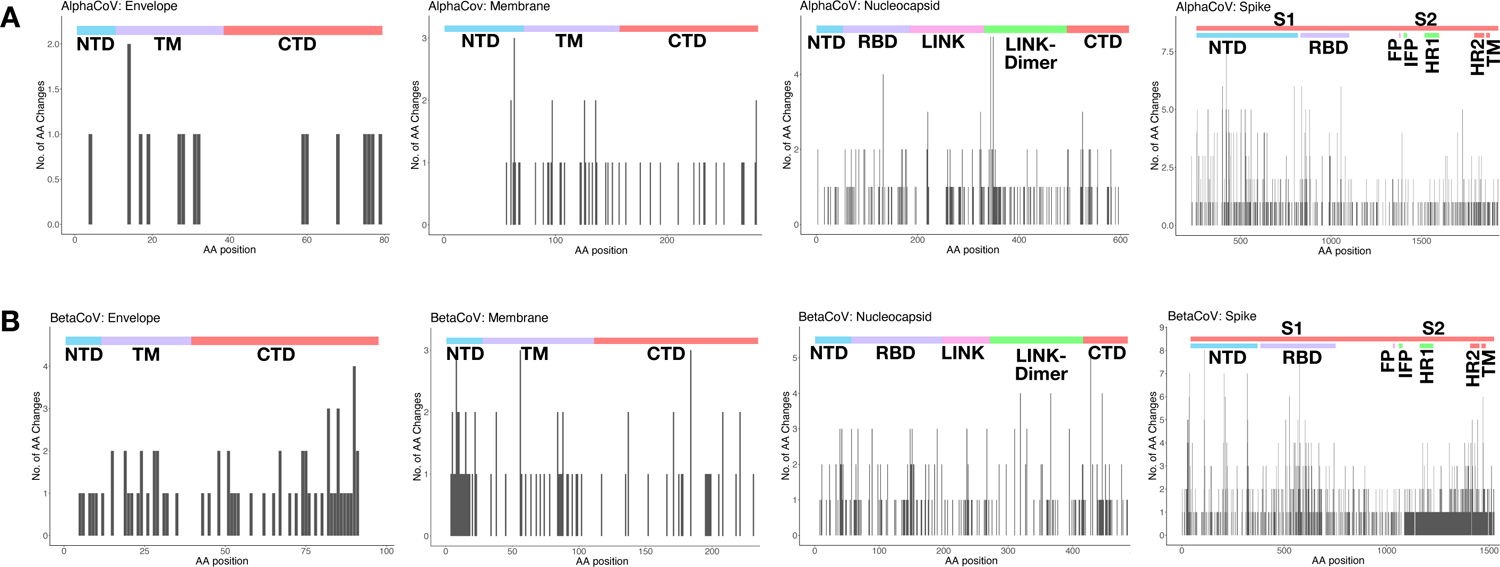
The number of inferred amino acid changes (AA) associated with the sHCoVs for AA positions in the envelope, membrane, nucleocapsid and spike proteins from datasets D4 and D5. Panel (A) represents the aggregated AA changes from the alphaCoVs 229E and NL63 while (B) represents the aggregated changes from the betaCoVs OC43 and HKU1. At the top of each plot, the functional domains or regions of the respective proteins are shown; NTD=N-terminal domain, TM=transmembrane domain, CTD=C-terminal domain, RBD=receptor binding domain, LINK=central linker domain, LINK-Dimer=dimerization domain, S1 subunit, S2 subunit, FP=fusion peptide, IFP=internal fusion peptide, HR1=heptad repeat 1, and HR2=heptad repeat 2.

The reconstructed AA changes were grouped by occurrence either along the host-jump branches or along the within-host branches in order to elucidate the nature of changes associated with the two evolutionary processes. The AA changes along the host-jump branches could highlight adaptive changes associated with host switches such as host receptor binding, while the AA changes along the within-host branches might highlight host-specific adaptive changes such as those that mediate immune escape.

The majority of the AA changes along the host-jump branches into humans were consistently associated with NL63 and HKU1 for all four proteins analyzed, individually accounting for 25-62% of the AA changes along these host-jump branches. However, for the within-human branches, HKU1 and OC43 were associated with the largest numbers of AA changes with the exception of the spike protein where 229E had the second highest number of AA changes after HKU1. It is important to note that the sHCoV species associated with the most AA changes along the host-jump and within-human branches had the longest branch lengths and the largest synonymous changes of all the sHCoVs, and therefore the higher number of AA changes does not necessarily reflect differences in selection.

For all four structural proteins, there was no single common AA position at which there were AA changes for all the four host-jump branches leading to the emergence of the sHCoV species. This might not be surprising considering that the four sHCoVs have different zoonotic origins (see section on <u>Zoonotic origins of the seasonal human coronaviruses</u>). However, the following positions had AA changes in the emergence of three sHCoV species: spike [152, 624, 680, 932, 1153, 1154, 1165, 1170], nucleocapsid [127], membrane [222], and envelope [27] (Table 5). Most of the shared positions in the sHCoV host-jump branches were also observed to have AA changes in 1-15 non-human to non-human host-jump branches and/or within-host branches in the MCC trees.

**Table 5:**
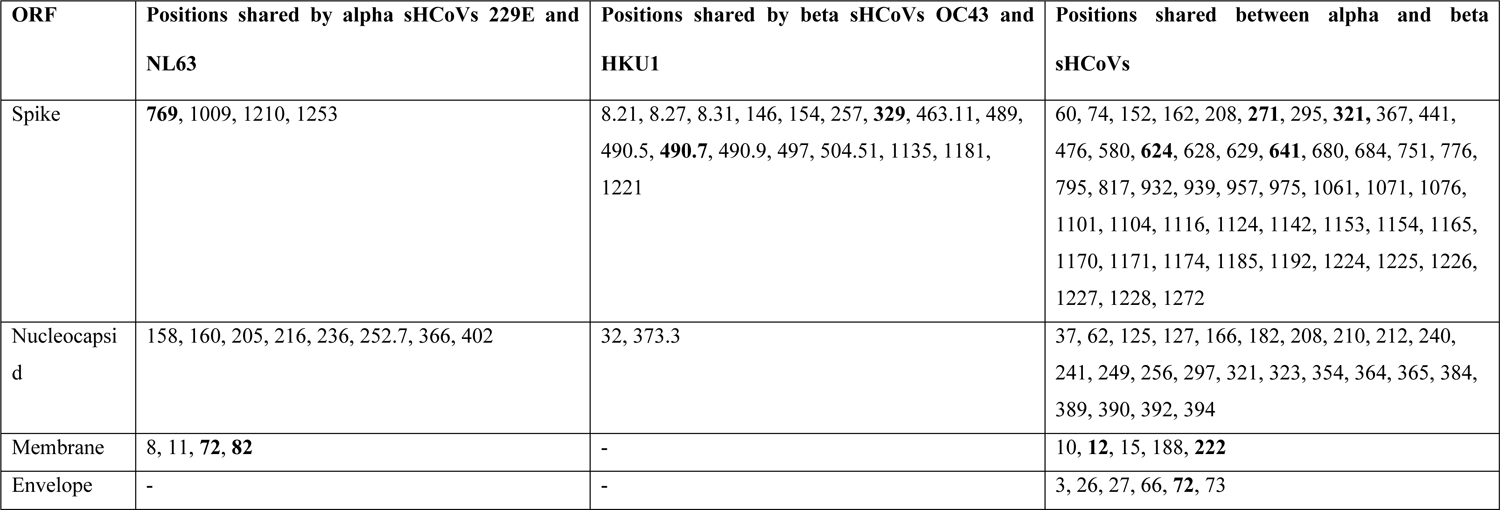
A list of positions with AA change in at least two host-jump branches leading to the sHCoVs. Where there was an AA insertion in the sHCoVs relative to Wuhan-Hu-1 SARS-CoV-2 reference genome, we used the X.Y positional notation where X is the reference genome position and Y is n^th^ sHCoV AA insertion. In bold are positions at which there were no AA changes in branches other than the host-jump branches leading to the sHCoVs.

While the majority of the AA changes at these common AA positions along the host-jump branches did not yield any obvious patterns, we broadly categorized some of the observed AA changes into (i) convergent changes to the same AA [e.g. NL63: L1224F and HKU1: G1224F], (ii) divergent changes from the same AA to different AAs [e.g. NL63: I769L and 229E: I769N], (iii) parallel AA changes [e.g. NL63: A1009S and 229E: A1009S], (iv) convergent AA changes with SARS-CoV-2 [e.g. OC43: N856K (Omicron VOC)], (v) completely reversed AA changes between sHCoV species [e.g. NL63: I1225V and HKU1: V1225I], and (vi) partially reversed AA changes between sHCoV species [e.g. NL63 S162Y and HKU1: N162S] (Supplementary Table 2).

The convergent AA changes across different host-jump branches for the sHCoVs comprised sHCoV pairs from the same genus or across genera. More than 80% of these convergent AAs along the host-jump branches for the four sHCoV structural proteins appeared fixed, i.e., there were no subsequent AA changes at these positions along the within-human branches. The parallel AA changes in the spike protein were only observed in sHCoVs from the same genus, and all but the N146K change were fixed.

The host-jump branches leading to sHCoVs NL63, HKU1 and OC43 experienced convergent AA changes with SARS-CoV-2 Wuhan-Hu-1 or a VOC in the spike protein, i.e. NL63: P80A [Beta VOC] [75], T375S [Wuhan-Hu-1] [76], HKU1: E339D [Omicron VOC] [77], F371L [Omicron VOC] [78], R969K [Omicron VOC], and OC43: N856K [Omicron VOC] [79]. For the nucleocapsid and membrane proteins, HKU1 [E63D] and NL63 [L82I], respectively, had convergent AA changes with Wuhan-Hu-1 CoV.

The partially or completely reversed changes for pairs of sHCoV host-jump branches might reflect different zoonotic origins or some other unknown adaptive characteristic but might also be non-adaptive changes that occurred at the same positions by chance. Most of the AA positions in the spike proteins with partially reversed AA changes between sHCoVs involved polar uncharged AAs (i.e., included serine, threonine, glutamine or asparagine). For HKU1 and OC43, however, these partially reversed AA changes in the spike protein all involved the positively charged AA lysine. Lastly, there were two completely reversed AA changes between HKU1 [D1165N and V1225I] and NL63 [N1165D and I1225V], and these changes were fixed in these two sHCoVs.

## Discussion

Using a multigene and a complete genomes approach, we find the evolutionary histories of sHCoVs to be more complex and uncertain than previously recognized, owing to frequent recombination of CoVs including within and between sHCoVs, which occurs at different rates. The origins of sHCoVs 229E and OC43 may depend on the gene studied. For HKU1, we propose the possibility of two independent non-human to human transmission events. We find shared AA substitutions along the non-human to sHCoV host-jump branches, even though these branches did not often appear to be under positive selection. Tracking the evolution of the sHCoVs will not only aid the understanding of their evolutionary history and the mechanisms for the potential emergence of novel human CoVs, but also has implications for development of diagnostics, vaccines and therapeutics.

The emergence of 229E from a bat CoV has been thought to be through camelids as intermediate hosts based on phylogenetic clustering, genetic similarity to, and serological detection of similar viruses in dromedary camels and alpacas [38–40]. In our analysis, the camelid CoV origins for 229E were not well supported, and instead independent transmissions of bat CoVs into humans and camelids appeared more likely. However, the possibility of reverse zoonosis of 229E into camelids would need further scrutiny as this would be in contrast with the MERS CoV whereby it is thought that humans represent an evolutionary dead-end for MERS-CoV [80].

For OC43, a less certain origin was observed than the previously proposed emergence from a bovine CoV; sHCoV OC43 clustered with ungulate and canine CoVs whose sequence data was not available at the time the bovine origin hypothesis was put forward [36]. We estimated bovine CoV to have 2-36% probability as the origin of the OC43, and for some of the ORFs, porcine, camel and murine CoVs had the highest probabilities for the origins of OC43. Our observations widen the repertoire of potential OC43 origin/intermediate hosts to include other ungulates, rabbits and canines; hosts that are either domesticated or wild but always in close proximity to humans.

Coronaviruses frequently recombine [18], and SARS-CoV-2 is already showing signs of recombination [81]. The process of recombination confers increased viral diversity and potential for inter-host transmission [82, 83]. We observed the sHCoVs to be recombinants of non-human CoVs, indicating that either the sHCoVs that emerged in humans were already recombinants of diverse non-human CoVs or the extant sHCoVs are a product of continuous viral back-and-forth transmission between human and non-human hosts. We report previously undocumented recombination within 229E, as well as recombination between the sHCoV species from the same genera. Our results demonstrate that recombination frequently occurs beyond the spike protein that is often the focus of recombination analyses. As viral coinfections are frequently identified in patients [3, 10], recombination between sHCoV species could be a real-world phenomenon that needs to be considered in studying the evolution of human CoVs in the SARS-CoV-2 era. To what extent these within and between sHCoV species recombination events facilitate the expansion of sHCoV variants and thereby the fitness of these sHCoVs warrants further investigation. Moreover, our observations raise questions about recombinant variants that might be missed by current sHCoV diagnostic panels.

Replication-associated mutations are a major source of genetic diversity in RNA viruses [84]. Recombination rates varied across the genome, between alpha and beta CoV genera, and between the sHCoVs. The higher rates of recombination in the spike protein relative to other parts of the genome has also been observed in other studies using different methods [85]. Two recent papers looking at rates of adaptation [86] and recombination [87] in sHCoVs reported that these two rates are concurrently elevated and decreased in different parts of the genome. The two studies not only corroborate our observations on recombination rate variation across the sHCoV genomes, but also highlight the important role of recombination in these CoVs’ adaptive processes considering that recombination was estimated to cause 0.8 times as many substitutions as point mutation events across the whole genome.

It was surprising that the MRCA of the two HKU1 genotypes analyzed collectively (HKU1_all) was dated much earlier than all the other sHCoVs, considering that HKU1 is the most recently isolated sHCoV [32]. A previous analysis using only the spike protein S1 domain of HKU1 had estimated MRCA dates in the 1950s [88].

Since the MRCAs of HKU1 A and B genotypes are herein dated more recently and a few years prior to the HKU1 first isolation date (similar to the lapse between the estimated MRCA and initial isolation dates for 229E and OC43), we propose two scenarios: (i) a single inter-host transmission into the human population with prolonged undetected circulation, or (ii) two independent inter-host transmission events into the human population of two closely related but very distinct viruses at similar time periods. Insufficient sampling is often characterized by large uncertainty (95% HPDs) in MRCA estimates [89] as was observed with HKU1. In addition, retrospective sampling has led to the identification of NL63 isolates such as the KF530110 (NL63/human/USA/838-9/1983) that are older than the first described NL63 isolate from 2005, and therefore it is not farfetched to think that HKU1 might have been in circulation in the human population without prior isolation and identification. Two independent transmission events at similar time periods could be supported by the consistently phylogenetically distinct HKU1 genotypes in multiple ORFs, in addition to the marked variation in genetic diversity and recombination rates. However, the best evidence for independent transmission of each HKU1 genotype into the human host would be phylogenetic clustering of each genotype with a murine CoV as a sister clade. Additionally, it would be worthwhile to investigate if the two HKU1 genotypes are characterized by distinct viral phenotypes such as clinical presentation and frequency of detection.

It is thought that the criteria that determines a virus’s inter-host transmission are availability of host receptor for virus binding and entry, permissiveness of the host cells to allow the virus to replicate, and avoidance of the host species-specific innate immune responses inhibiting viral replication [90]. Therefore, a virus that has undergone a cross-species transmission will leave some adaptive footprint in its amino acid profile. Ancestral reconstruction of the sHCoV spike, nucleocapsid, membrane and envelope protein sequences revealed that at least 16% and up to 57% of the AA positions in these proteins had at least one AA substitution across the genealogy of the sHCoVs. Of the four structural proteins, the spike protein had the largest number of one amino acid substitutions, which is expected because it is the longest of these four proteins and also has a more significant role in the adaptation of CoVs to new hosts [91]. However, except for the host-jump branches leading to NL63, HKU1_all and HKU1 genotype B for the nucleocapsid, the inter-host transmission process leading to the emergence of the sHCoVs did not appear to be under positive selection. Noteworthily, these host-jump branches may not be under adaptation as HKU1 and NL63 had the longest host-jump and within-host branches, largest synonymous changes, and in some cases had unrealistically high ω values. A recent computational study of the spike protein inferred adaptive evolution in 229E and OC43 viruses and none at all in HKU1 and NL63 [86]. Taken together, it could be that inter-host transmission of CoVs from a non-human to a human host, as well as continued circulation in the human population, can occur even without detectable positive selective pressure that is often expected to drive the optimization of a virus to its host [92].

## Conclusions

As a consequence of the CoVs’ ability to recombine, mutate, and infect multiple hosts, SARS-CoV-2 is certainly neither the last CoV to spill-over into the human population nor the last with pandemic potential. As more human and non-human viral genome sequences become available, there is an opportunity and necessity to further investigate the evolutionary histories of CoVs to achieve a better understanding of how these viruses jump between species and establish recurrent infections. These investigations could lead to the design of control strategies. Continuous surveillance of non-human CoV hosts would be essential for early identification of potential zoonotic outbreaks, especially in largely under-sampled areas such as Africa whose bat CoVs appeared ancestral to the sHCoVs 229E and NL63 [39,40,93].

## Acknowledgments

We thank Darren Martin at the University of Cape Town (SA), Maciej Boni at the Pennsylvania State University (USA), and Xavier Didelot at the University of Warwick (UK) for discussions and suggestions on recombination in viruses. Also thanking Philippe Lemey at KU Leuven (Belgium) for suggestions on BEAST analyses. The opinions expressed in this article are those of the authors and do not reflect the view of the National Institutes of Health, the Department of Health and Human Services, or the United States government.

## Author Contributions

Conceptualization: JRO, MIN, NST

Methodology: JRO, JLC, DJS, MIN, NST

Investigation: JRO, JLC

Visualization: JRO

Writing - original draft: JRO, NST

Writing - review & editing: JRO, JLC, DJS, MIN, NST

## Data and Materials Availability

No new data were generated or analyzed in support of this research. The XML-format files containing the data and model parametrization will be made available on GitHub (url will be added upon revision).

---

**Supplementary Figure 1:**
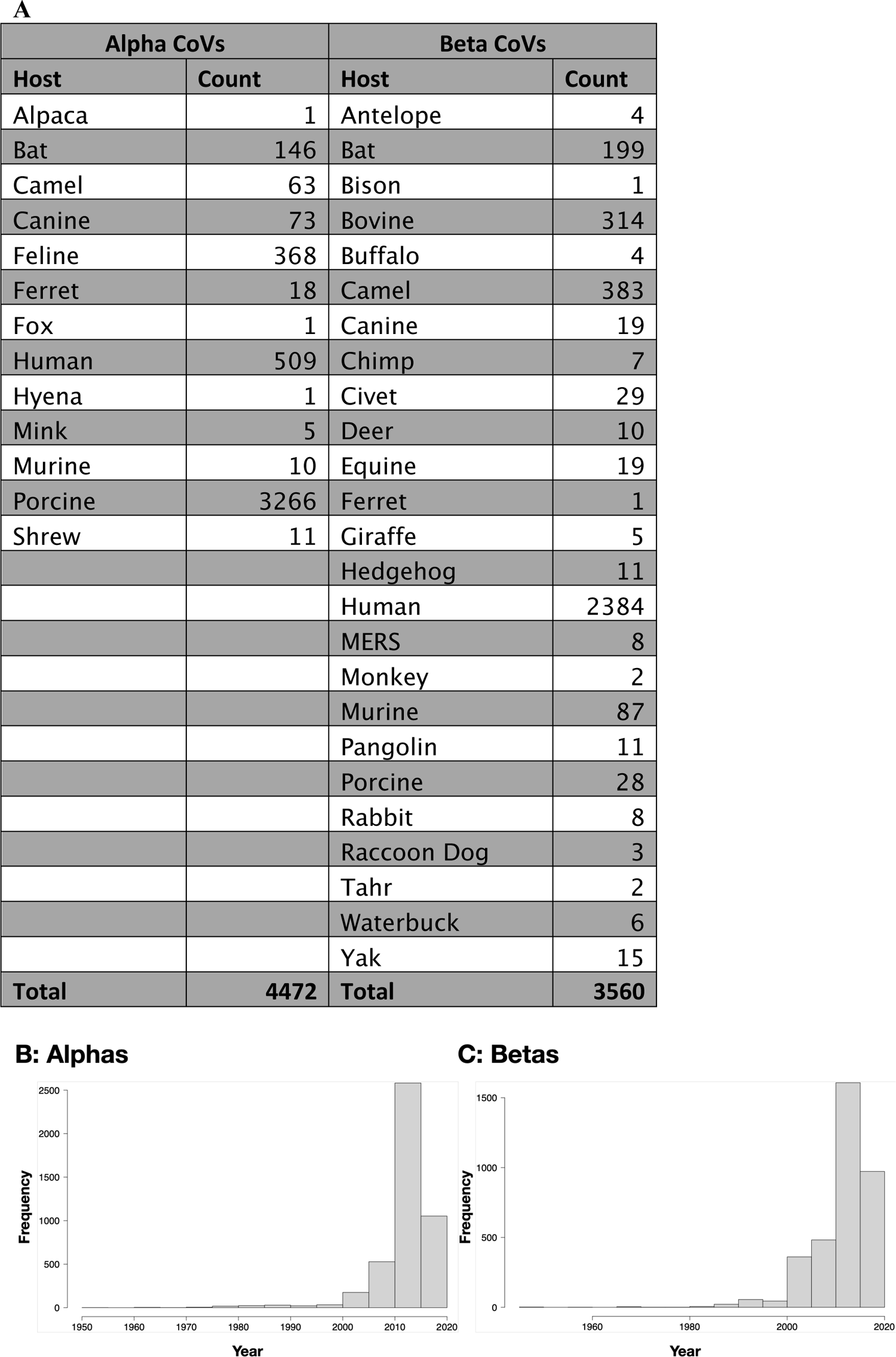
Downloaded and cleaned dataset D1, by host (A) and year (B-C).

**Supplementary Figure 2:**
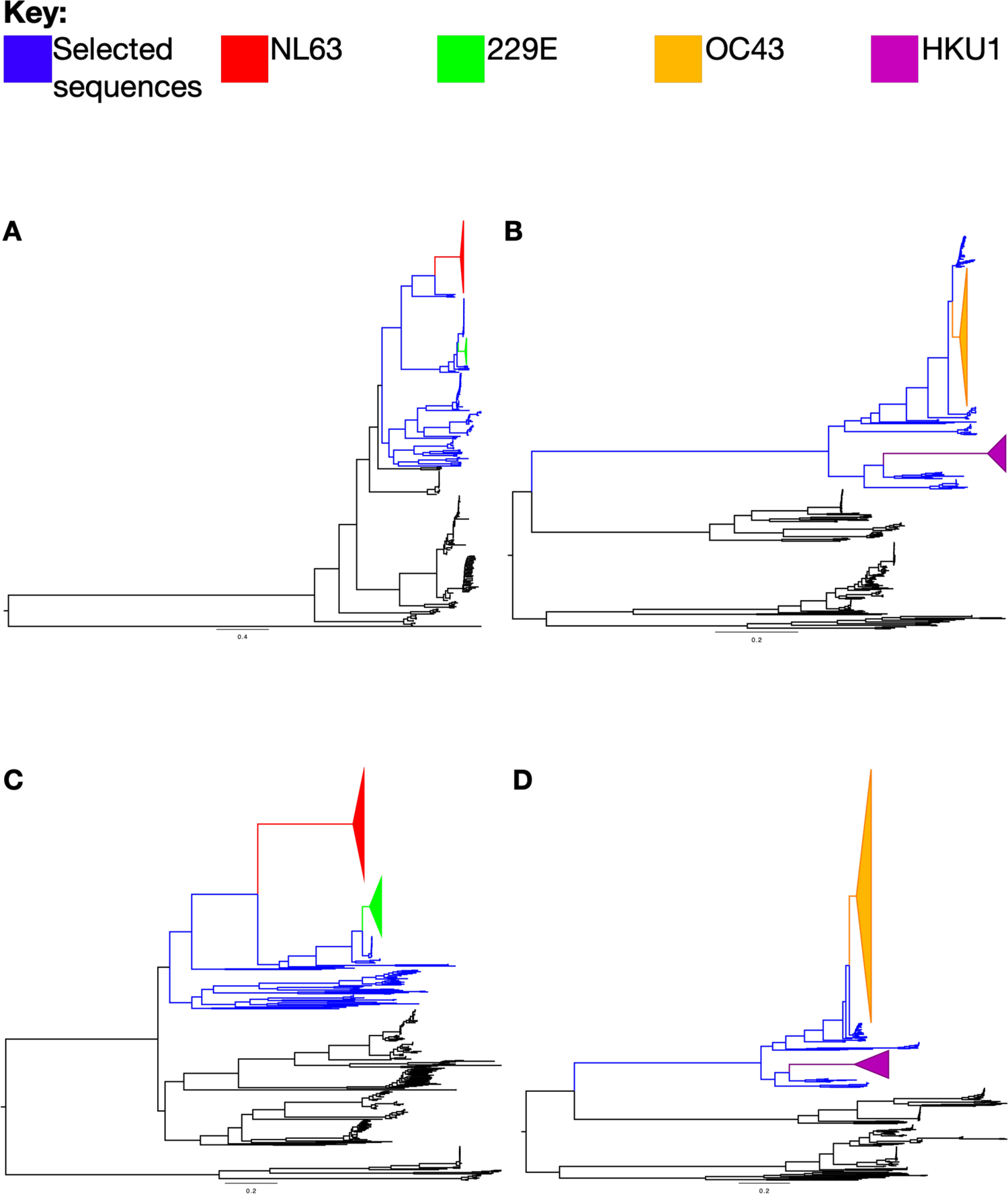
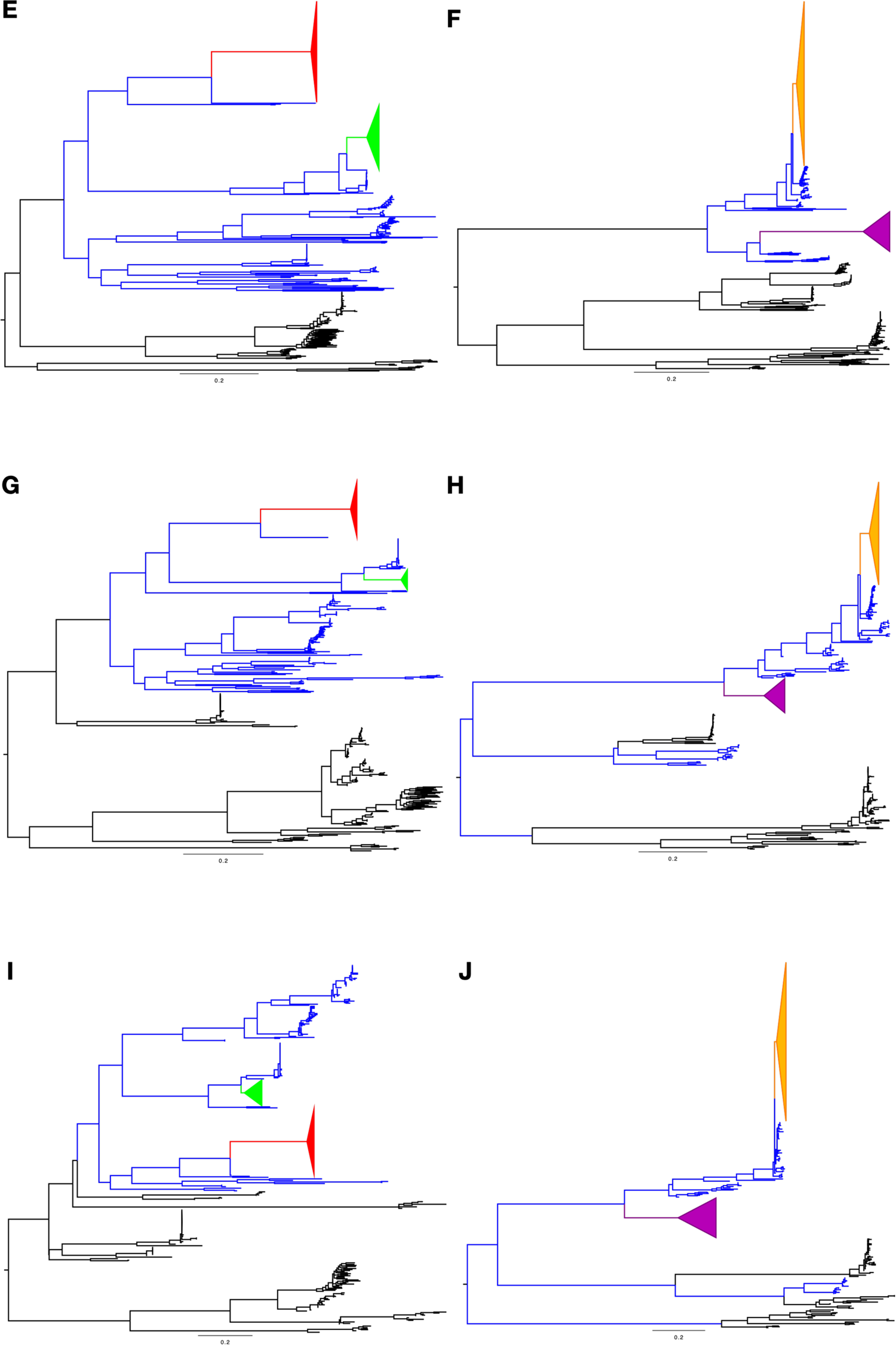
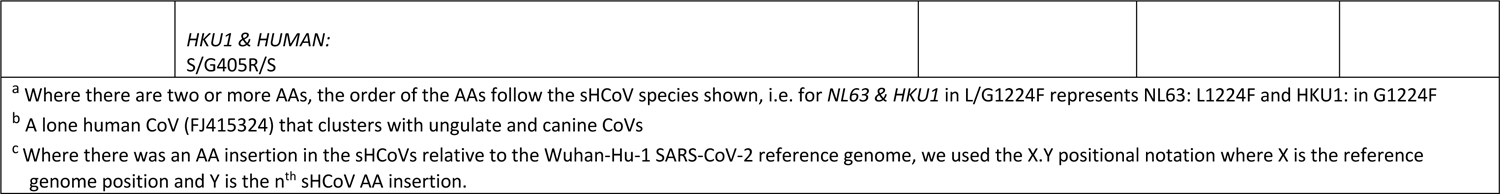
Maximum Likelihood (ML) phylogenetic trees from dataset D2 for whole genomes (A-B), and the Spike (C-D), Nucleocapsid (E-F), Membrane (G-H) and Envelope (A-J) proteins. Branches highlighted in blue represent the non-sHCoV sequences selected for the final analyses in addition to the seasonal human coronaviruses (sHCoVs) colored in red (NL63), green (229E), orange (OC43) and purple (HKU1). On these trees, CoVs from hosts other than the sHCoVs and those that share a MRCA as per the ML trees had been subsampled to ∼ 30-40 sequences per host-clade, i.e. if there were two porcine clades positioned on different parts of a tree then they were subsampled independently.

**Supplementary Figure 3:**
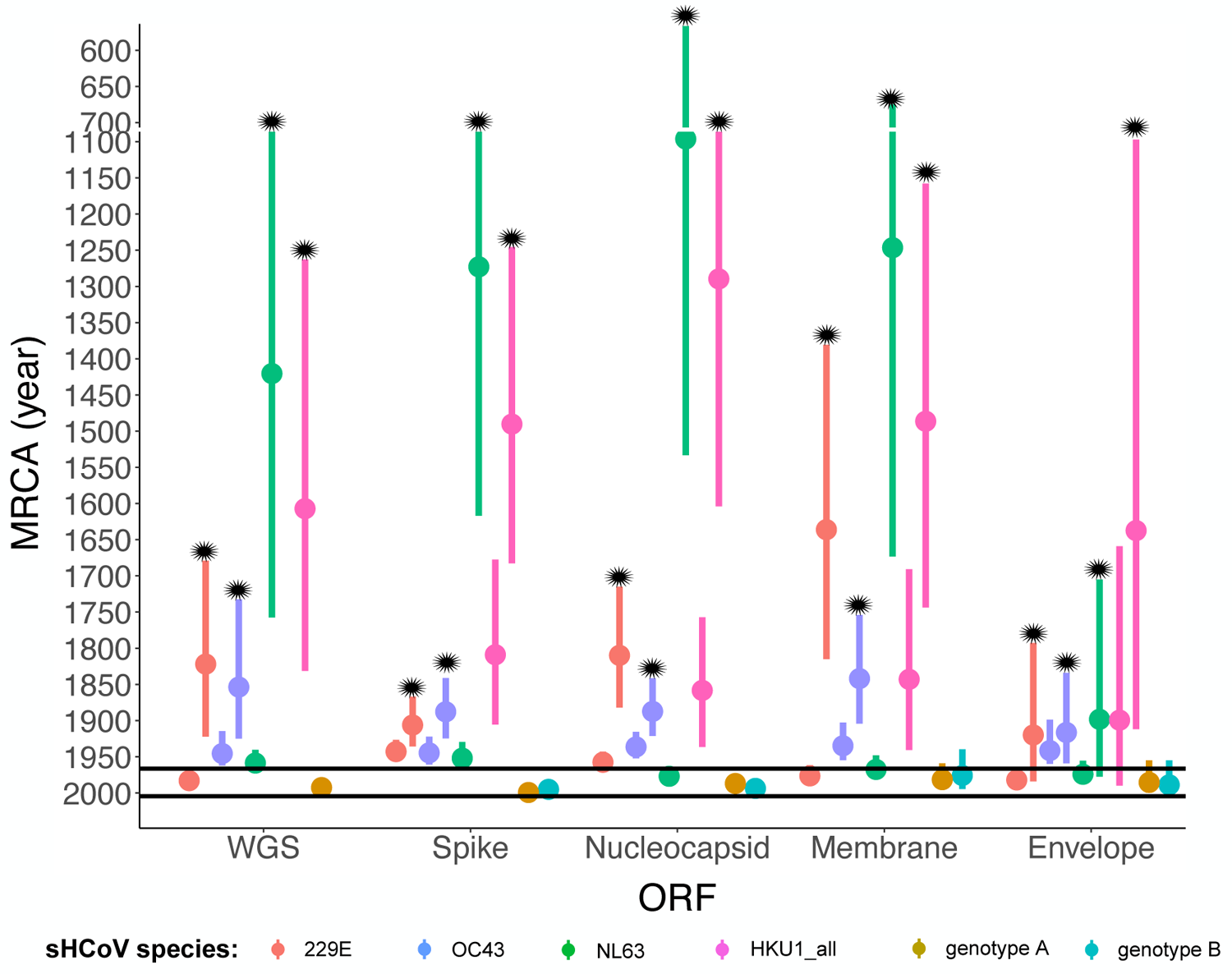
Estimates of the MRCA age for full genomes and four open reading frames (dataset D5) of the seasonal human coronavirus species. The black horizontal lines represent the dates of first isolation for 229E (1966), OC43 (1967), NL63 (2004) and HKU1 (2005). The star (*) symbol shows the parents to the MRCAs of the sHCoVs. The WGS is missing data points for HKU1_all (collective for both genotypes) and genotype B as sequences for genotype B were all removed in the generation of recombination-free WGS dataset D5.

**Supplementary Figure 4:**
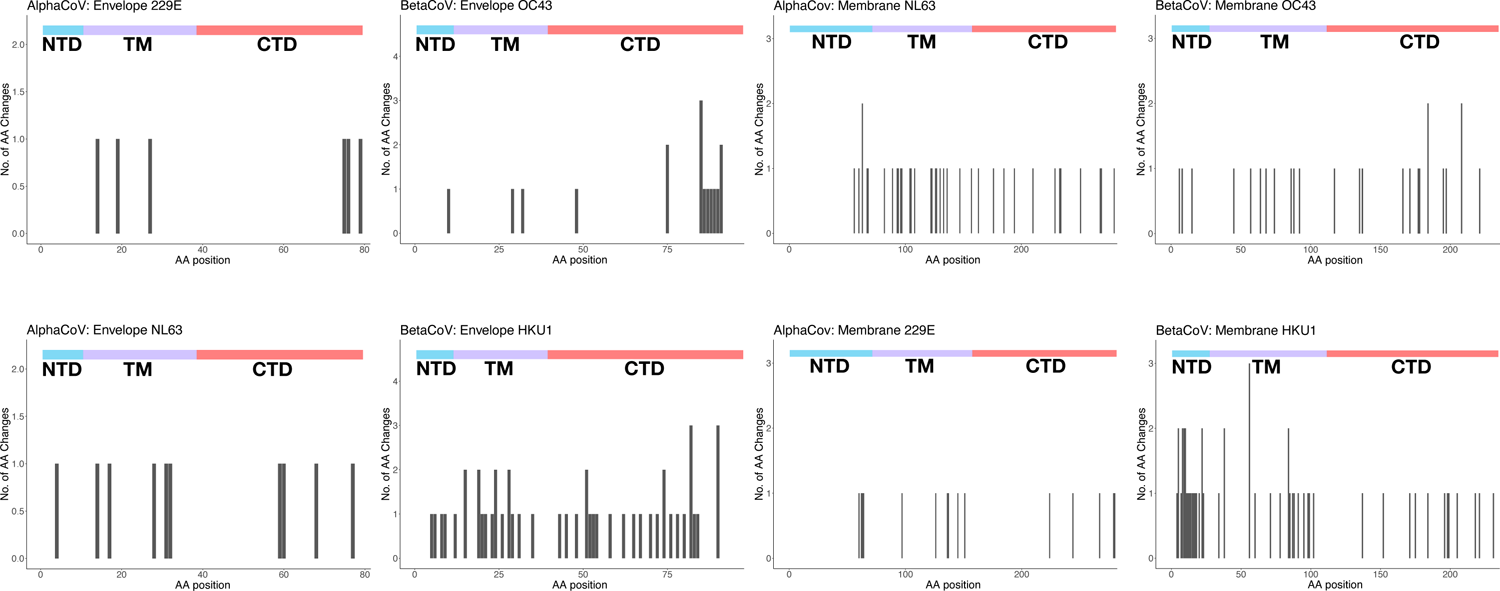
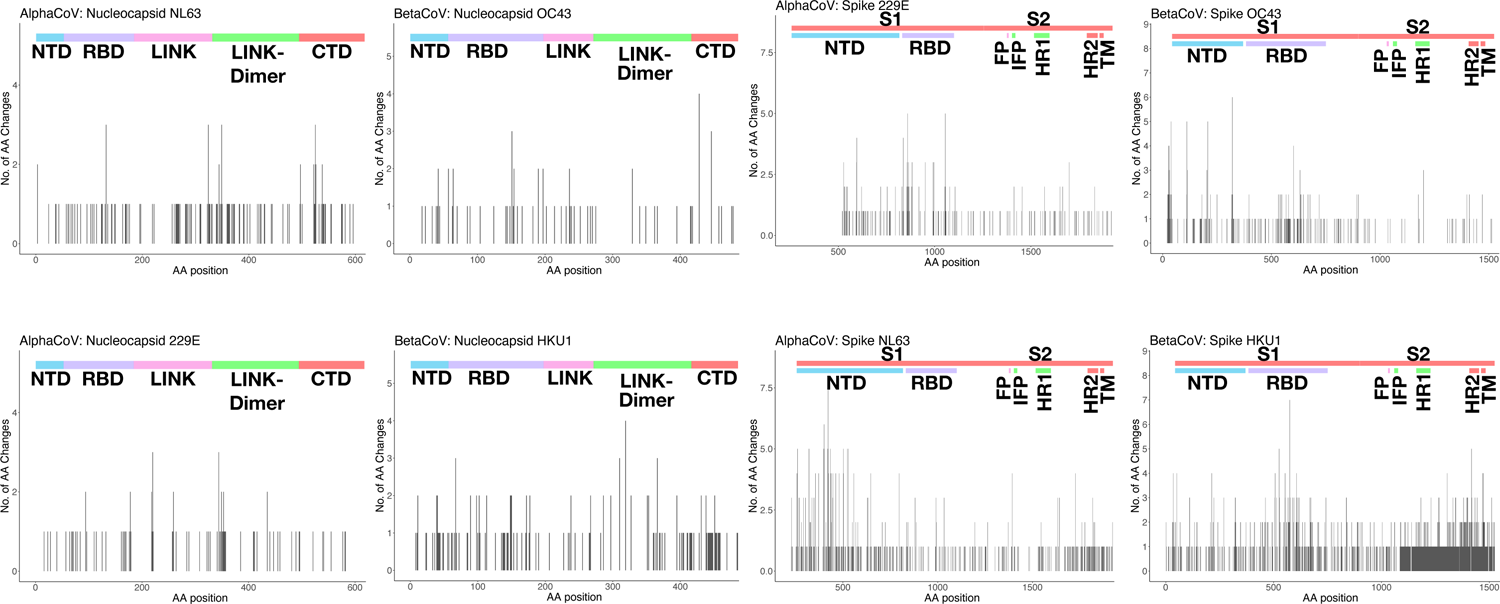
The number of inferred amino acid changes (AA) within sHCoV clades for AA positions in the envelope, membrane, nucleocapsid and spike proteins from datasets D4 and D5. At the top of each plot, the functional domains or regions of the respective proteins are shown; NTD=N-terminal domain, TM=transmembrane domain, CTD=C-terminal domain, RBD=receptor binding domain, LINK=central linker domain, LINK-Dimer=dimerization domain, S1 subunit, S2 subunit, FP=fusion peptide, IFP=internal fusion peptide, HR1=heptad repeat 1, and HR2=heptad repeat 2.

**Supplementary Table 1:**
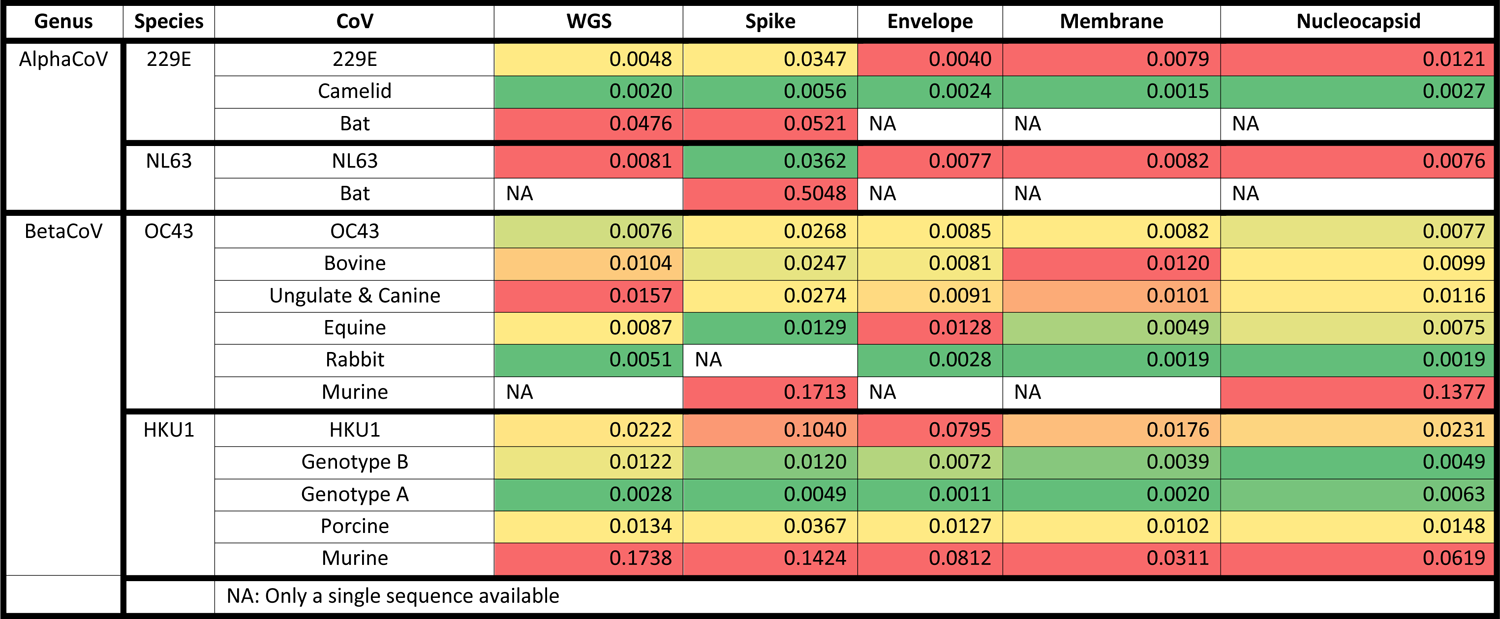
Mean pairwise genetic distances for various CoV host-clades. The CoV host-clade sequences selected for this analysis were based on ML tree topologies in *Supplementary Figure 1*, selecting host-clades that were sister clades to the sHCoVs. However, when the isolates that shared a recent ancestor with humans, rather than one in the more distant past, were represented by a single sequence, the mean pairwise genetic distance could not be calculated on the single sequence. We also included CoVs from hosts not closely related to the sHCoVs for comparison. For each sHCoV species and ORF/WGS, the cells are colored from the highest (red) to the lowest (green) genetic distance.

**Supplementary Table 2:**
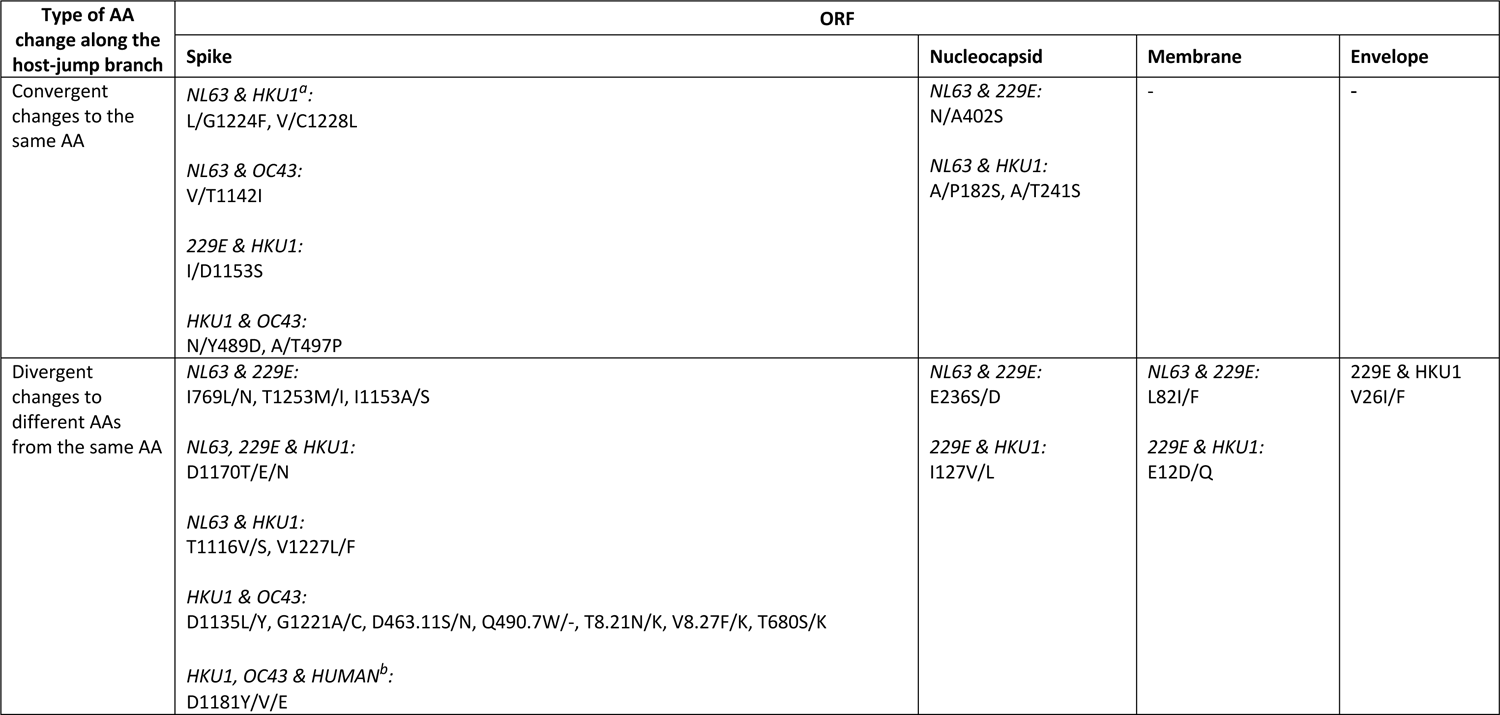
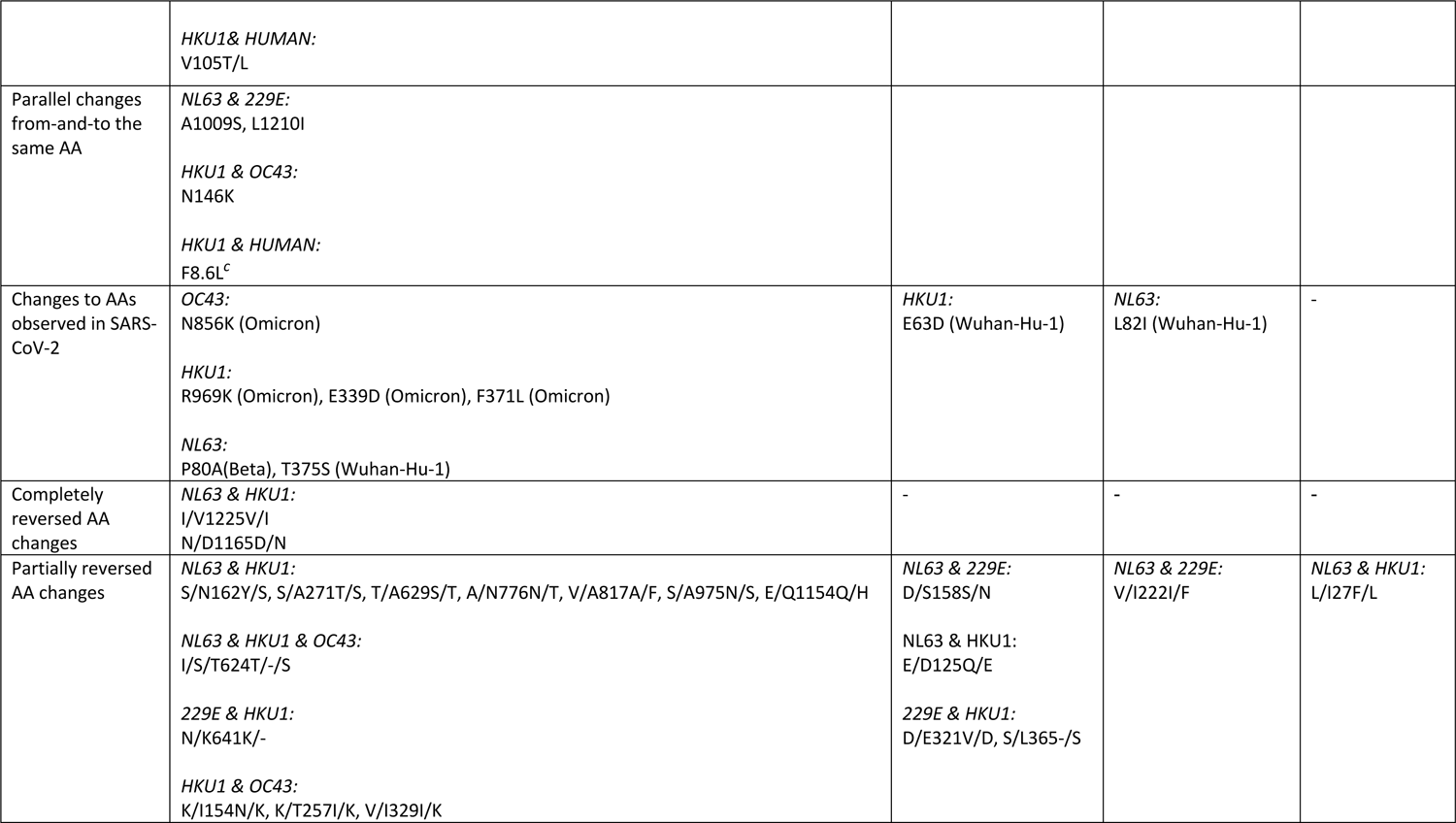
A select amino acid changes occurring along the host-jump branches leading to the emergence of the sHCoVs

## Supplementary Files Legends

Supplementary File 1: <u>subsampling_data.xlsx</u>

Supplementary File 2: <u>MCC Trees</u>

Supplementary File 3: RDP Recombination Analysis

Supplementary File 4: AllAAChanges.xlsx

